# Neuronal Dysfunction and Isotretinoin-Mediated Modulation in Patient-Derived iPSC Models of Smith–Magenis Syndrome: Parallels with Sustained Clinical Improvement in a Patient

**DOI:** 10.64898/2026.11.28.769862

**Authors:** Idan Rosh, Yara Hussein, Aviram Shemen, Ritu Nayak, Utkarsh Tripathi, Hagit Sadis, Diogo Cordeiro, Tali Garin-Shkolnik, Ilana Schlesinger, Shani Stern

## Abstract

Smith–Magenis syndrome (SMS) is a rare neurodevelopmental disorder characterized by intellectual disability, behavioral abnormalities, autistic-like features, sleep disturbances, and distinctive physical characteristics. SMS is primarily caused by a heterozygous deletion of chromosome 17p11.2 encompassing the *Retinoic Acid Induced 1* (*RAI1*) gene or by pathogenic variants within *RAI1*. Although *RAI1* haploinsufficiency is considered the principal molecular cause of SMS, the mechanisms underlying neuronal dysfunction and clinical variability are not fully understood, and the development of targeted therapeutic strategies remains an unmet clinical need. Given the involvement of *RAI1* in transcriptional regulation and its reported association with retinoid signaling, we investigated the potential of isotretinoin (13-cis-retinoic acid), a vitamin A derivative, to modulate disease-associated neuronal phenotypes.

Human induced pluripotent stem cells (hiPSCs) derived from two individuals with SMS and mutation-negative familial controls were differentiated into midbrain dopaminergic (DA) and hippocampal dentate gyrus (DG) neurons. Neuronal function was evaluated using whole-cell patch-clamp electrophysiology at different maturation stages, while transcriptional responses to 1 µM isotretinoin were characterized using bulk RNA sequencing.

Whole-cell patch-clamp recordings revealed distinct patient-specific electrophysiological abnormalities, with neuronal excitability and excitatory synaptic activity deviating from control levels. Isotretinoin treatment partially attenuated these abnormalities, shifting selected electrophysiological parameters toward control values by increasing evoked firing in hypoexcitable neurons and reducing hyperexcitability in patient-derived DA neurons. However, treatment responses varied across patients, neuronal populations, and maturation stages, with limited improvement observed in hyperexcitable hippocampal neurons.

Transcriptomic analysis revealed extensive isotretinoin-associated changes involving synaptic signaling, extracellular matrix organization, neuronal development, and retinoid metabolism. In DA neurons, 5,475 genes were differentially expressed following treatment. Hippocampal neurons similarly exhibited retinoid-responsive transcriptional changes, although the magnitude and functional consequences differed between patient lines.

Longitudinal clinical observations in an individual with SMS and early-onset Parkinson’s disease (PD) revealed sustained improvements in motor function, balance, postural stability, and selected numerical abilities following the initiation of oral isotretinoin treatment, most notably the recovery of independent ambulation. Together, these findings demonstrate substantial heterogeneity in SMS-associated neuronal dysfunction and identify isotretinoin as a modulator of neuronal excitability and transcriptional regulation. The experimental findings, together with the longitudinal clinical observations, provide a rationale for further investigation of retinoid-based therapeutic strategies in SMS.

## Introduction

SMS is a rare neurodevelopmental disorder characterized by a distinctive combination of cognitive, behavioral, neurological, and physical abnormalities. SMS is most commonly caused by a heterozygous interstitial deletion of chromosome 17p11.2 encompassing the *RAI1* gene, although pathogenic variants affecting *RAI1* can also produce the syndrome in the absence of a chromosomal deletion [1–4]. The estimated prevalence of SMS is approximately 1 in 15,000–25,000 individuals, although underdiagnosis may contribute to uncertainty in this estimate. Although SMS is characterized by a recognizable clinical phenotype, substantial interindividual variability exists in the manifestation and severity of its associated features. [3, 5–7].

The clinical manifestations of SMS include intellectual disability, developmental and speech delays, sleep disturbances, and characteristic behavioral abnormalities, including impulsivity, aggression, repetitive behaviors, and self-injury [5, 7–9]. Cognitive and adaptive functioning vary considerably among affected individuals, with difficulties frequently observed in communication, daily living skills, and motor development [10–12]. Some clinical features of SMS overlap with those observed in autism spectrum disorder (ASD), suggesting potential similarities in the neurodevelopmental pathways implicated in these conditions. However, the cellular and molecular mechanisms contributing to neuronal dysfunction and clinical variability in SMS remain unclear.

The RAI1 gene encodes a large nuclear protein implicated in transcriptional regulation and neuronal development. The canonical human RAI1 protein comprises 1,906 amino acids, with the majority of its coding sequence concentrated in exon 3, a region in which numerous pathogenic variants associated with SMS have been identified[13–15]. The RAI1 protein contains several characteristic structural features, including an N-terminal polyglutamine tract, two polyserine-rich regions, putative nuclear localization signals, and a C-terminal plant homeodomain (PHD) finger, consistent with its proposed involvement in chromatin-associated transcriptional regulation [13, 16].

Experimental studies have implicated RAI1 in the regulation of neuronal gene expression, circadian rhythms, and metabolic homeostasis [17–19]. In particular, Huang et al. demonstrated that Rai1 is involved in molecular and neural processes relevant to SMS-associated behavioral abnormalities [19]. Subsequent work established a role for RAI1 in activity-dependent nascent transcription and homeostatic synaptic scaling, providing a mechanistic link between RAI1 deficiency, altered neuronal gene regulation, and synaptic dysfunction [20]. Together, these findings suggest that reduced RAI1 expression may disrupt multiple cellular processes required for normal neuronal development and function.

hiPSC technology provides an opportunity to investigate the cellular consequences of disease-associated genetic alterations in patient-derived neuronal populations. By retaining the donor-specific genetic background, iPSC-based models can reveal disease-associated abnormalities in neuronal development, excitability, and synaptic function [21]. Directed differentiation into specific neuronal subtypes further enables the investigation of cell-type-dependent vulnerability and disease mechanisms. Human iPSC PD models revealed synaptic deficits in DA neurons derived from the patients compared to healthy controls [22–24]. ASD models similarly reported synaptic deficits in cortical and hippocampal neurons [25–27] that first started as neuronal hyperexcitability at an early age that later developed into synaptic deficits at 2-3 months after the start of the differentiation.

Midbrain DA neurons and DG neurons represent two functionally distinct neuronal populations relevant to motor, cognitive, and behavioral regulation. Midbrain DA neurons contribute to motor control, motivation, and reward-related processes, whereas hippocampal DG neurons play important roles in learning, memory, and pattern separation. Established differentiation approaches permit the generation of these neuronal populations from human pluripotent stem cells, providing experimentally accessible systems for investigating disease-related alterations in neuronal physiology [28, 29]. Examining both neuronal populations may therefore help determine whether the consequences of *RAI1* haploinsufficiency are shared across neuronal subtypes or exhibit cell-type-specific characteristics.

Despite advances in understanding the genetic basis of SMS, therapeutic approaches capable of addressing its underlying molecular and neuronal abnormalities remain limited. A study demonstrated that adeno-associated virus-mediated CRISPR activation (rAAV-CRISPRa) could increase expression of the remaining functional *Rai1* allele and rescue selected disease-associated phenotypes in an SMS mouse model [30]. These findings provide proof of concept for approaches aimed at compensating for *RAI1* haploinsufficiency, although the extent to which restoration of RAI1 expression can normalize diverse neuronal phenotypes remains uncertain.

Retinoic acid (RA), a biologically active derivative of vitamin A, regulates gene expression through retinoic acid receptor-dependent signaling and plays important roles in nervous system development, neuronal differentiation, and maturation [31, 32]. In addition to its established roles in neuronal development, RA signaling is also involved in the regulation of synaptic strength and neuronal homeostasis [33, 34]. This functional overlap with processes affected by RAI1 deficiency provides a rationale for investigating retinoid signaling as a potential modulator of neuronal dysfunction in SMS.

Iisotretinoin (13-cis-retinoic acid) is a synthetic retinoid with established clinical applications, including its use as maintenance therapy in the treatment of high-risk neuroblastoma [35]. Previous investigations have also examined its effects on hippocampus-dependent learning and neurogenesis, highlighting the potential influence of retinoid exposure on neuronal function [36, 37]. Although *RAI1* was originally identified through studies demonstrating retinoic acid-induced expression of its mouse homolog in P19 embryonal carcinoma cells[2, 15, 38], this observation does not establish that isotretinoin directly restores *RAI1* expression or reverses neuronal abnormalities associated with *RAI1* haploinsufficiency. Therefore, the potential effects of isotretinoin on SMS-associated neuronal dysfunction require direct experimental investigation.

In the present study, we investigated the cellular and molecular consequences of *RAI1* haploinsufficiency using hiPSC-derived midbrain DA and DG neurons generated from two individuals with SMS and two unaffected first-degree relative controls. One of the patients (56 years of age) presented with early onset PD [39]and this was the motivation to differentiate DA neurons in addition to hippocampal neurons. We characterized neuronal excitability and synaptic activity using whole-cell patch-clamp recordings and examined transcriptional alterations through bulk RNA sequencing. We further evaluated whether isotretinoin treatment could modulate disease-associated electrophysiological and transcriptional abnormalities across distinct neuronal populations and patient genetic backgrounds. In parallel, longitudinal clinical observations from one participant with SMS who independently received isotretinoin for a dermatological indication provided exploratory clinical context for the experimental findings. Together, these approaches aimed to characterize patient-specific neuronal dysfunction in SMS and investigate the potential of isotretinoin to modify disease-associated cellular phenotypes.

## Materials and Methods

### Ethics statement

The study was approved by the Institutional Review Board (IRB) of the University of Haifa (project identification code: 282-22; April 27, 2023). Written informed consent was obtained from all participants or their respective legal guardians prior to participation in the study. No financial compensation or other incentives were provided to the participants

### Peripheral blood mononuclear cell isolation

Biological samples used to establish patient-derived hiPSC lines were collected before the initiation of isotretinoin treatment. The patient subsequently began receiving oral isotretinoin in April 2023 as part of routine clinical care, following a recommendation of the clinician due to dermatology issues. Experimental investigations involving patient-derived neuronal cultures were initiated several months after the commencement of clinical treatment. Thus, the original patient samples were obtained before systemic isotretinoin exposure, and the subsequent in vitro experiments were conducted independently of the initial clinical treatment decision. This experimental framework enabled the investigation of isotretinoin-associated cellular and molecular responses in patient-derived neurons without prior in vivo isotretinoin exposure of the sampled cells.

Peripheral blood samples were collected from four participants using standard venipuncture procedures. The study cohort comprised two male individuals diagnosed with SMS, each carrying a pathogenic genetic alteration affecting RAI1, and two unaffected first-degree relatives serving as genetically related controls.

Peripheral blood was collected into 8 mL BD Vacutainer Cell Preparation Tubes (CPT; BD Biosciences, 362780) and processed immediately following collection. Samples were centrifuged at 1,800 × *g* for 30 min at 23°C using a swinging-bucket rotor. Following centrifugation, the plasma layer was carefully removed, and peripheral blood mononuclear cells (PBMCs) were collected from the mononuclear cell layer above the gel barrier. The isolated cells were washed with calcium- and magnesium-free Dulbecco’s phosphate-buffered saline (DPBS; Sartorius, 02-023-1A) and centrifuged again at 300 × *g* for 15 min.

Following centrifugation, the supernatant was discarded, and the cell pellet was resuspended in calcium- and magnesium-free DPBS for cell counting. The PBMCs were subsequently resuspended in cryopreservation medium consisting of 90% fetal bovine serum (FBS; sterile- filtered; Merck, F7524-500ML) and 10% dimethyl sulfoxide (DMSO; Thermo Scientific, 127791000). Approximately 2 × 10 cells were aliquoted into individual cryovials. The cryovials were placed in a pre-cooled Styrofoam container and stored at −80°C overnight before being transferred to liquid nitrogen for long-term storage

### Clinical Assessment and Longitudinal Follow-up

Longitudinal clinical follow-up was conducted in a patient with SMS carrying an *RAI1* mutation who developed progressive parkinsonian manifestations beginning in 2020[39]. The clinical follow-up was coordinated by the research team in collaboration with the treating neurologist, who performed neurological examinations and documented the patient’s clinical status. Assessments primarily consisted of clinical neurological examinations and functional observations rather than a predefined battery of standardized neurological or neuropsychological tests.

Clinical evaluations included observations of gait, balance, tremor, postural control, ocular movements, and functional motor abilities, including independent walking, running, squatting, and standing on one leg. Selected cognitive abilities were assessed through clinical observations of sequential counting, simple arithmetic, attention, and conversational performance. Findings were documented across successive clinical visits, and available neurological reports were reviewed to characterize longitudinal changes.

The patient had been receiving levodopa/carbidopa (Dopicar) for approximately two years before the initiation of isotretinoin treatment and continued receiving it throughout the follow-up period. Oral isotretinoin (Curatane) was introduced in April 2023 at a dosage of 10 mg three times weekly. The dosage was increased to 20 mg three times weekly on July 13, 2023, and reduced to 10 mg three times weekly on November 28, 2023, following the identification of mild abnormalities in liver-related laboratory parameters. Treatment tolerability was monitored through clinical follow-up and available laboratory assessments.

### Reprogramming of PBMCs into iPSCs Using Sendai Virus-Mediated Gene Transfer

PBMCs were thawed four days before reprogramming and cultured in 24-well plates containing StemPro™ SFM medium (Thermo Fisher Scientific, 10639011) supplemented with 100 ng/mL stem cell factor (SCF; PeproTech, 300-07), 100 ng/mL FMS-like tyrosine kinase 3 ligand (FLT3L; PeproTech, 300-19), 20 ng/mL interleukin-3 (IL-3; PeproTech, 200-03), and 20 ng/mL interleukin-6 (IL-6; PeproTech, 200-06). During this period, half of the culture medium (50%) was replaced daily.

On the day of reprogramming (day 0), approximately 5 × 10 PBMCs were transduced using the CytoTune-iPS Sendai Reprogramming Kit (Life Technologies, A16518). Following transduction, cells were centrifuged at 1,160 × *g* for 30 min at 35°C, seeded into 24-well plates, and incubated overnight at 37°C in a humidified atmosphere containing 5% CO . The following day, cells were washed to remove residual extracellular viral particles and transferred to low-attachment 12-well plates. After two days, cells were transferred onto Matrigel-coated plates and maintained with medium changes every two days.

Beginning on day 7, the culture medium was gradually replaced with ReproTeSR medium (STEMCELL Technologies, 05296), with complete transition achieved by day 8. Emerging iPSC colonies were observed between days 15 and 21, manually selected, and transferred onto fresh Matrigel-coated plates for further expansion in mTeSR medium (STEMCELL Technologies, 100-0276).

Once iPSC colonies reached an appropriate size, individual colonies were manually picked and transferred onto Matrigel-coated six-well plates for further expansion. Pluripotency was confirmed by immunocytochemical staining for the pluripotency markers TRA-1-60, SOX2, and OCT4.

The generated iPSC lines were assigned the following abbreviated designations to facilitate their identification throughout the study, including in electrophysiological and transcriptomic analyses. The SMS patient-derived lines UOHi009 and UOHi055 were designated T and Z, respectively. Their corresponding family-matched control lines were UOHi008 (S), derived from the unaffected sister of patient T, and UOHi056 (ZC), derived from the unaffected mother of patient Z.

### Differentiation of hiPSCs into DG Granule Neurons

To generate DG granule neurons, we followed a previously published differentiation strategy [21, 29]. hiPSCs lines of 2 patients and respective controls were expanded to approximately 40% confluency prior to differentiation. Embryoid bodies (EBs) were generated by mechanically dissociating colonies using 1 U/mL Dispase (STEMCELL Technologies Technologies, 07923) and transferring them to low-attachment culture plates containing mTeSR™ Plus medium supplemented with 10 μM Y-27632 ROCK inhibitor (Tocris Bioscience, 1254).

EBs were cultured for 20 days in an anti-caudalizing medium consisting of DMEM/F12, 1% GlutaMAX™, 2% B27 Supplement minus Vitamin A (Gibco, 12587-010), 1% N2 Supplement (Gibco, 17502-048), 100 nM LDN-193189 (Biogems, 1066208), 5 μM XAV939 (Biogems, 2848932), 1 μM cyclopamine (Biogems, 4445185), and 10 μM SB431542 (Biogems, 3014193). Subsequently, EBs were plated onto poly-L-ornithine/laminin-coated culture dishes in DMEM/F12 supplemented with GlutaMAX™, N2 Supplement, B27 Supplement minus Vitamin A, and laminin.

After one-week, neural rosettes were manually selected based on their characteristic morphology, dissociated using Accutase (Innovative Cell Technologies, AT104), and replated onto poly-L-ornithine (Sigma, P3655)/laminin (Sigma, L2020)-coated plates. Cells were maintained in neural progenitor cell (NPC) medium composed of DMEM/F12, GlutaMAX™, B27 Supplement minus Vitamin A, N2 Supplement, laminin, and 20 ng/mL FGF2.

Neuronal differentiation was induced by supplementing the culture medium with 200 nM ascorbic acid (Biogems, 5088177), 1 mM dibutyryl-cyclic adenosine monophosphate (cAMP; Adooq, A15914-5), 1 μg/mL laminin, 20 ng/mL brain-derived neurotrophic factor (BDNF; PeproTech, AF450-02), and 10 ng/mL Wnt3a for 14 days. Three weeks after the initiation of differentiation, the basal medium was replaced with BrainPhys™ medium (STEMCELL Technologies, 05790) to promote neuronal maturation.

### Differentiation of hiPSCs into midbrain DA neurons

To generate in vitro midbrain DA neurons, we followed a previously published differentiation protocol[28, 40, 41]. Briefly, patient- and control-derived hiPSCs were dissociated into a single-cell suspension using Accutase, replated onto Matrigel-coated culture plates in mTeSR™ Plus medium, and allowed to proliferate prior to the initiation of differentiation at approximately 50% confluency (designated as day 0). Cells were gradually transitioned from KSR medium, consisting of DMEM/F12 supplemented with 1% GlutaMAX™, 25% KnockOut™ Serum Replacement (KO-SR), 1% non-essential amino acids (NEAA), and 0.1 mM β-mercaptoethanol, to N2 medium composed of DMEM/F12, GlutaMAX™, and 1% N2 Supplement between days 5 and 10. On day 11, cultures were switched to B27 medium consisting of Neurobasal™ medium(Gibco, 21103049) supplemented with 1% GlutaMAX™, 2% B27 Supplement, 10 ng/mL BDNF, 10 ng/mL glial cell line-derived neurotrophic factor (GDNF), 1 ng/mL transforming growth factor-β3 (TGF- β3), 0.2 mM ascorbic acid, and 0.5 mM cAMP.

Throughout the differentiation process, patterning factors and small molecules were added to the cultures as follows: 10 μM SB431542 from days 0–4; 100 nM LDN-193189 from days 0– 12; 2 μM Purmorphamine (Biogems, 4831086), 0.25 μM Smoothened agonist (SAG; Biogems, 9128694), and 100 ng/mL FGF8b (PeproTech, 100-25) from days 1–6; and 3 μM CHIR99021 (Biogems, 2520691) from days 5–12.

Between days 20 and 25, neuronal cultures were dissociated and replated, and subsequently maintained in B27 medium until day 30. At this stage, the basal medium was replaced with BrainPhys™ medium to promote neuronal maturation and synaptic connectivity. Samples for bulk RNA sequencing were collected on differentiation day 45 for DA neurons and day 35 for DG neurons. Whole-cell patch-clamp electrophysiological recordings were performed at weeks 6 and 10 for DA neurons and weeks 4 and 8 for DG neurons.

Retinoid supplementation differed between the neuronal differentiation protocols. DA neurons were maintained in medium containing standard B27 Supplement from day 11 of differentiation, whereas DG neurons were cultured with B27 Supplement without Vitamin A throughout differentiation and maturation. Within each neuronal population, untreated and isotretinoin-treated cultures were maintained under otherwise matched differentiation conditions.

### Whole-cell patch-clamp electrophysiology

Whole-cell patch-clamp recordings were performed on DG granule neurons and midbrain DA neurons differentiated from hiPSC lines derived from SMS patients and healthy controls. Given that hiPSC-derived neurons exhibit distinct maturation timelines compared with primary mouse neurons, electrophysiological recordings were conducted at developmental stages selected according to previously established protocols [21, 23, 40]. Specifically, recordings were performed at two time points following the initiation of neuronal differentiation: 4 and 8 weeks for DG granule neurons, and 6 and 10 weeks for DA neurons.

To investigate the effects of isotretinoin on neuronal development and electrophysiological function, SMS patient-derived cultures were treated with 1 µM isotretinoin (13-cis-retinoic acid) beginning on day 1 of neuronal differentiation. Isotretinoin was replenished at each medium change throughout the differentiation period until electrophysiological assessment. Untreated patient-derived cultures and healthy control cultures were maintained in parallel. Electrophysiological experiments were conducted across three independent differentiation batches for each cell line, with consistent findings observed across batches.

For electrophysiological recordings, coverslips containing neuronal cultures were transferred to a recording chamber perfused with HEPES-buffered artificial cerebrospinal fluid (ACSF) containing (in mM): 139 NaCl (Sigma-Aldrich, S9888), 4 KCl (Sigma-Aldrich, P3911), 2 CaCl (Sigma-Aldrich, 223506), 1 MgCl (Merck, 7791-18-6), 10 HEPES (Sigma-Aldrich, H0887), and 10 D-glucose (Sigma-Aldrich, G7021). The ACSF was adjusted to pH 7.5 and an osmolality of 310 mOsm/kg, as measured using a Vapro 5600 vapor pressure osmometer (Wescor, USA).

Patch pipettes were fabricated from borosilicate glass capillaries (Sutter Instrument, BF150-75-10) using a P-1000 micropipette puller (Sutter Instrument, Novato, CA, USA) and fire-polished to achieve a final resistance of 9–12 MΩ. Pipettes were filled with an intracellular solution containing (in mM): 130 K-gluconate, 6 KCl, 4 NaCl, 10 Na-HEPES, 0.2 K-EGTA, 0.3 GTP, 2 Mg-ATP, 0.2 cAMP, and 10 D-glucose. The intracellular solution was adjusted to pH 7.5 and an osmolality of 290–300 mOsm/kg.

All recordings were performed at room temperature using a MultiClamp 700B amplifier (Molecular Devices, San Jose, CA, USA) connected to a Digidata 1550B digitizer (Molecular Devices, San Jose, CA, USA) and controlled using MultiClamp 700B Commander and pCLAMP 11 software. Signals were acquired at a sampling rate of 20 kHz and analyzed using Clampfit 10 (Molecular Devices) and MATLAB R2014b (MathWorks, Natick, MA, USA).

### Analysis of Electrophysiological Recordings

Electrophysiological data were analyzed using custom-written MATLAB scripts based on previously described methods [21].

#### Voltage-gated sodium and potassium currents

Neurons were voltage-clamped at −60 mV, and 400-ms voltage steps were applied over a voltage range of −100 to +90 mV. Voltage-gated sodium (Na ) currents were quantified as the difference between the peak inward current and the stabilized current measured following channel inactivation. Fast potassium (K ) currents were quantified as the peak outward currents occurring within the first few milliseconds following depolarization, whereas slow K currents were measured at the end of the 400-ms depolarizing step. Current amplitudes were normalized to the membrane capacitance of each cell to account for differences in cell size. Membrane capacitance was determined using the capacitance measurement function in Clampex software. Statistical comparisons between groups were performed using one-way analysis of variance (ANOVA).

#### Evoked action potentials

Neuronal excitability was assessed in current-clamp mode, with the membrane potential maintained at −60 mV by applying a constant holding current. Depolarizing current injections of 400 ms duration were delivered in 3 pA increments, beginning 12 pA below the steady holding current. A total of 38 current steps were applied to each neuron. The total number of evoked action potentials (APs) was calculated as the cumulative number of action potentials generated across all 38 current steps. The maximum number of evoked APs was defined as the highest number recorded during any single current step. Frequency–current (F–I) relationships were generated by calculating the firing frequency at each current injection step, defined as the number of action potentials divided by the 400-ms stimulus duration. Mean firing frequencies ± standard error of the mean (SEM) were plotted as a function of injected current.

#### Synaptic current analysis

Excitatory postsynaptic currents (EPSCs) were recorded in voltage-clamp mode at a holding potential of −60 mV. For each neuron exhibiting synaptic activity, the mean EPSC amplitude and corresponding standard error (SE) were calculated. Cumulative amplitude distributions were generated for each experimental group and condition. EPSC frequency was calculated for each neuron by dividing the total number of detected synaptic events by the recording duration. Neurons without detectable synaptic activity were assigned an EPSC frequency of 0 Hz. Mean EPSC frequencies ± SE were subsequently calculated for each experimental group. For evoked AP and EPSC measurements, group differences were assessed primarily using Kruskal–Wallis tests, with Mood’s median or permutation tests where indicated in the figures, followed by Holm– Bonferroni-adjusted pairwise comparisons where applicable.

#### Spontaneous action potentials

Spontaneous neuronal firing was assessed in current-clamp mode without depolarizing current injection, with the membrane potential maintained at approximately −45 mV. Spontaneous action potentials were recorded over a 60-s period, and the spontaneous firing rate was calculated as the number of action potentials per recording period. Statistical comparisons between groups were performed using nonparametric tests.

Immunocytochemistry (ICC)

Cells cultured on coverslips were fixed with 4% paraformaldehyde for 15 min and subsequently washed three times with DPBS for 5 min each. Cells were then permeabilized and blocked in PBS containing 0.1–0.2% Triton X-100 and 10% horse serum. Following blocking, coverslips were incubated overnight at 4°C with primary antibodies diluted in blocking solution.

For midbrain DA neurons, the following primary antibodies were used: anti-tyrosine hydroxylase (TH; Abcam, ab129991; 1:500), anti-MAP2 (Abcam, ab92434; 1:500), and anti-cleaved caspase-3 (Cell Signaling Technology, 9664; 1:500). For DG neurons, primary antibodies included anti-PROX1 (Abcam, ab199359; 1:4000), anti-MAP2 (Abcam, ab92434; 1:500), and anti-cleaved caspase-3 (Cell Signaling Technology, 9664; 1:500).

The following day, coverslips were washed with DPBS and incubated for 60 min at room temperature with DAPI (Abcam, ab228549; 1:2500) and the appropriate fluorophore-conjugated secondary antibodies. Coverslips were then washed three times with DPBS, mounted onto glass slides using Fluoromount-G mounting medium (SouthernBiotech, 0100-01), and allowed to dry overnight while protected from light.

Fluorescence imaging was performed using a Leica THUNDER Imager and a ZEISS LSM 980 confocal microscope equipped with Airyscan 2. Acquired images were analyzed using ImageJ and Imaris 9.8 (Oxford Instruments) software.

### Bulk RNA Sequencing and Bioinformatic Analysis

#### RNA isolation, library preparation, and sequencing

Bulk RNA sequencing (RNA-seq) was performed on hiPSC-derived DA and DG neurons generated from patients with SMS carrying *RAI1* mutations and mutation-negative controls. Approximately 3–5 × 10 neurons were collected per sample. DA and hippocampal neurons were harvested on differentiation days 45 and 35, respectively. Cells were lysed using TRIzol reagent, and total RNA was subsequently purified using the Zymo RNA Clean & Concentrator Kit according to the manufacturer’s instructions.

For each experimental condition, two RNA-seq samples were obtained from separate neuronal cultures. Sequencing libraries were prepared using the TruSeq Stranded mRNA Library Prep Kit (Illumina), followed by paired-end sequencing (2 × 150 bp) on the Illumina NovaSeq X platform, targeting approximately 40 million reads per sample.

#### Sequencing quality control, alignment, and gene quantification

Raw sequencing reads were assessed for quality using FastQC within the Galaxy platform. Reads were aligned to the human reference genome GRCh38 (hg38) using STAR (v2.7.11b), with two-pass mapping disabled and a splice-junction overhang of 100 nucleotides. Uniquely mapped reads were assigned a mapping quality (MAPQ) score of 60.

Gene-level read counts were generated using HTSeq-count (v2.1.2), applying reverse-stranded counting and a minimum alignment quality threshold of 10.

#### Differential gene expression analysis

Downstream bioinformatic analyses were performed in R using custom scripts, with DA and hippocampal neuronal datasets analyzed separately. Genes with at least 10 counts in two or more samples were retained for differential expression analysis using DESeq2. Differentially expressed genes (DEGs) were defined by a Benjamini–Hochberg-adjusted *P* value < 0.05 and an absolute log fold change > 0.5.

For the DA dataset, differential expression was evaluated between untreated SMS patient-derived neurons and mutation-negative controls, as well as between isotretinoin-treated (ISO; 1 µM) and untreated patient-derived neurons. Comparisons with the familial and unrelated mutation-negative controls were performed separately. Positive log fold changes represented higher expression in the first condition of each comparison.

For the hippocampal dataset, differential expression between SMS patient-derived neurons and mutation-negative familial controls was assessed using a family-adjusted DESeq2 design (∼ family + status). Separate analyses compared untreated patient-derived neurons with their corresponding familial controls and isotretinoin-treated patient-derived neurons with familial controls. Family membership was included as a covariate to account for baseline transcriptional differences between the two families, while the patient-versus-control contrast estimated a common SMS-associated expression effect across families.

To evaluate the combined transcriptional response to isotretinoin in hippocampal neurons, only patient-derived samples were included in a separate DESeq2 model (∼ line + treatment), with patient-line identity incorporated as a covariate to account for baseline transcriptional differences between donors. Additional patient-specific treatment comparisons were performed independently for lines T and Z using the design ∼ treatment. For these analyses, positive log fold changes indicated higher expression in isotretinoin-treated neurons relative to untreated cultures.

Isotretinoin treatment was restricted to patient-derived neuronal cultures; mutation-negative controls were maintained without isotretinoin treatment.

#### Principal component analysis and data visualization

Principal component analysis (PCA) was performed using variance-stabilized expression values generated with DESeq2. The 1,000 genes exhibiting the highest expression variability across samples were selected for PCA, or all available genes when fewer than 1,000 were present.

Differential expression results were visualized using volcano plots displaying log fold changes against −log -adjusted *P* values. Selected genes associated with neuronal identity, synaptic function, retinoid signaling, and neuroinflammation were further examined using expression heatmaps and normalized expression plots.

#### Functional enrichment analysis

Functional enrichment analysis was performed using the R package clusterProfiler. Gene Ontology (GO) enrichment was evaluated across Biological Process (BP), Molecular Function (MF), and Cellular Component (CC) categories, together with Kyoto Encyclopedia of Genes and Genomes (KEGG) pathway enrichment.

Ensembl gene identifiers were mapped to the identifiers required for the corresponding enrichment analyses using org.Hs.eg.db. Enrichment was assessed for the complete set of DEGs and separately for upregulated and downregulated genes, where applicable. Statistical significance was determined using Benjamini–Hochberg-adjusted *P* values < 0.05.

#### Immune-associated transcriptional analysis

Immune- and inflammation-associated transcriptional changes were evaluated by targeted examination of annotated chemokine, cytokine, NF-κB-associated, interferon-responsive, complement, and glial-associated genes in the differential-expression tables. The same differential-expression criteria used in the primary analyses (Benjamini–Hochberg-adjusted P < 0.05 and |log fold change| > 0.5) were applied. Significant GO Biological Process terms related to immune and inflammatory signaling were identified from the existing over- representation analyses. These exploratory analyses describe transcript abundance and GO annotations, rather than direct measurements of inflammatory pathway activity. Patient- specific contrasts were interpreted independently; no formal neuronal-type-by-treatment interaction test was performed.

#### Biological replication and interpretation

For each hippocampal SMS patient-derived line, untreated and isotretinoin-treated neuronal cultures originated from the same differentiation experiment, with two RNA-seq samples collected per condition. Similarly, the DA RNA-seq samples represented separate cultures originating from a single differentiation batch.

## Results

### Clinical Observations and Longitudinal Follow-up

Patient T, an individual with SMS with a pathogenic genetic alteration affecting RAI1, developed progressive parkinsonian manifestations beginning in 2020 [39], including tremor, recurrent falls, impaired balance, and progressive gait difficulties. Dopaminergic therapy with carbidopa/levodopa (Dopicar, 25/250 mg) was initiated in 2021 at a dosage of one-quarter tablet three times daily, resulting in partial improvement in tremor and fall frequency. The dosage was subsequently increased to one-half tablet three times daily. Nevertheless, substantial motor impairment persisted, and the patient remained unable to ambulate independently, requiring physical assistance. Neurological examination before isotretinoin treatment also revealed restricted upward gaze, impaired eye contact, and difficulties performing basic numerical tasks. In April 2023, oral isotretinoin (Curatane) was introduced at a dosage of 10 mg three times weekly due to persistent skin problems that included acne and dryness, while the existing dopaminergic regimen was maintained.

During subsequent neurological follow-up, significant improvements in motor function, balance, postural control, and ocular movements were observed. Approximately one year after isotretinoin initiation, the patient was able to walk and run independently, with additional improvements in the ability to jump, squat, stand on one leg without support, and maintain an upright posture. Tremor was no longer clinically apparent during later examinations, although assistance remained necessary when navigating stairs for an additional year. Improvements were also observed in selected numerical tasks. Whereas the patient initially demonstrated difficulties counting sequentially beyond 10, subsequent examinations documented independent counting to 27 and, on some occasions, higher numbers, alongside the ability to perform simple arithmetic operations. However, performance varied between assessments, attention and concentration difficulties persisted, and although no clear improvement in verbal abilities was observed, the patient demonstrated improved ability to maintain eye contact during conversations and appeared more attentive and engaged, with an improved ability to listen, comprehend, and respond appropriately.

On July 13, 2023, the isotretinoin dosage was increased to 20 mg three times weekly. Following the detection of mild abnormalities in liver-related laboratory parameters, the dosage was reduced to 10 mg three times weekly on November 28, 2023. Subsequent laboratory assessments demonstrated normalization of the previously abnormal parameters. The patient continues receiving isotretinoin at the reduced dosage alongside levodopa/carbidopa through November 2026, with independent ambulation reportedly maintained. Although the temporal association between isotretinoin initiation and the observed neurological improvements is noteworthy, the observational nature of this single-patient case, concomitant dopaminergic treatment, and potential contributions of other clinical factors preclude definitive attribution of these improvements to isotretinoin.

Patient T-derived DA neurons exhibit reduced excitability and excitatory postsynaptic activity, with selective improvement following isotretinoin treatment

Whole-cell patch-clamp recordings were performed to assess neuronal excitability and excitatory synaptic activity in iPSC-derived DA neurons from SMS patients T and Z, healthy first-degree relatives as controls, and isotretinoin-treated patient-derived cultures. Electrophysiological properties were evaluated at weeks 6 and 10 of differentiation. Patient-derived cultures were exposed to 1 µM isotretinoin beginning on the first day of differentiation, with treatment maintained throughout neuronal differentiation. Recordings were obtained from three independent differentiation batches per cell line. Neuronal excitability was assessed by quantifying the total and maximum numbers of evoked APs, while excitatory synaptic activity was evaluated by measuring EPSC frequency and amplitude.

At week 6, untreated DA neurons derived from patient T exhibited significantly reduced excitability compared with healthy controls, as indicated by lower total and maximum numbers of evoked APs (Figure 1a–e; n = 71 control and 69 patient neurons; both P < 0.001). Isotretinoin treatment significantly increased both parameters relative to untreated patient-derived neurons (n = 19 treated neurons; both P < 0.001). Representative membrane potential recordings illustrated the reduced firing capacity of untreated patient-derived neurons and the increased evoked firing observed following treatment (Figure 1c–e).

**Figure 1.**
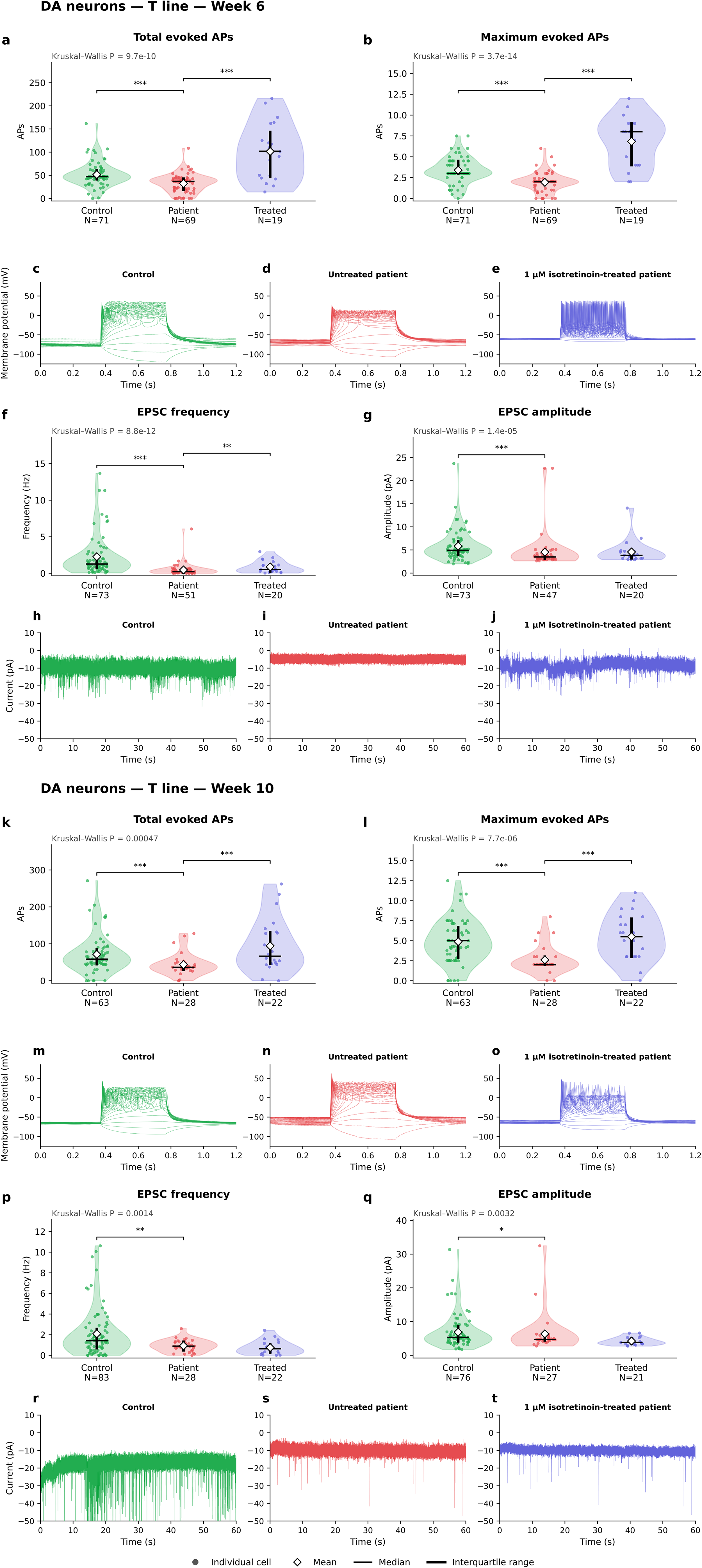
Electrophysiological characterization of SMS patient-derived DA neurons and the effects of isotretinoin treatment at weeks 6 and 10 of differentiation. Whole-cell patch-clamp recordings were performed in hiPSC-derived DA neurons from a healthy control and a patient with SMS (line T). Patient-derived neurons were either untreated or exposed to 1 µM isotretinoin beginning on the first day of differentiation and maintained throughout neuronal differentiation. **Week 6 (a–j): (a, b)** Quantification of the total (a) and maximum (b) numbers of evoked APs in control (n = 71), untreated SMS (n = 69), and isotretinoin-treated SMS (n = 19) neurons. Untreated patient-derived neurons exhibited significantly reduced total and maximum numbers of evoked APs compared with controls (both P < 0.001), whereas isotretinoin treatment significantly increased both parameters relative to untreated patient-derived neurons (both P < 0.001). **(c–e)** Representative current-clamp recordings showing evoked AP responses to sequential depolarizing current injections in control (c), untreated SMS (d), and isotretinoin-treated SMS (e) neurons. **(f, g)** Quantification of EPSC frequency (f) and amplitude (g). EPSC frequency was assessed in control (n = 73), untreated SMS (n = 51), and isotretinoin-treated SMS (n = 20) neurons, whereas EPSC amplitude was assessed in control (n = 73), untreated SMS (n = 47), and isotretinoin-treated SMS (n = 20) neurons. Untreated patient-derived neurons exhibited significantly reduced EPSC frequency and amplitude compared with controls (both P < 0.001). Isotretinoin treatment significantly increased EPSC frequency relative to untreated patient-derived neurons (P < 0.01), whereas EPSC amplitude was not significantly altered by treatment (P > 0.05). **(h–j)** Representative voltage-clamp recordings illustrating spontaneous EPSCs in control (h), untreated SMS (i), and isotretinoin-treated SMS (j) neurons. **Week 10 (k–t): (k, l)** Quantification of the total (k) and maximum (l) numbers of evoked APs in control (n = 63), untreated SMS (n = 28), and isotretinoin-treated SMS (n = 22) neurons. Untreated patient-derived neurons exhibited significantly reduced total and maximum numbers of evoked APs compared with controls (both P < 0.001), whereas isotretinoin treatment significantly increased both parameters relative to untreated patient-derived neurons (both P < 0.001). **(m–o)** Representative current-clamp recordings showing evoked AP responses to sequential depolarizing current injections in control (m), untreated SMS (n), and isotretinoin-treated SMS (o) neurons. **(p, q)** Quantification of EPSC frequency (p) and amplitude (q). EPSC frequency was assessed in control (n = 83), untreated SMS (n = 28), and isotretinoin-treated SMS (n = 22) neurons, whereas EPSC amplitude was assessed in control (n = 76), untreated SMS (n = 27), and isotretinoin-treated SMS (n = 21) neurons. Untreated patient-derived neurons exhibited significantly reduced EPSC frequency (P < 0.01) and amplitude (P < 0.05) compared with controls. Isotretinoin treatment did not significantly alter either EPSC frequency or amplitude relative to untreated patient-derived neurons (both P > 0.05). **(r–t)** Representative voltage-clamp recordings illustrating spontaneous EPSCs in control (r), untreated SMS (s), and isotretinoin-treated SMS (t) neurons. Violin plots show distributions of individual neuronal measurements; points represent individual neurons, white diamonds indicate means, horizontal black lines indicate medians, and vertical black bars indicate interquartile ranges. Sample sizes (n) are indicated for each group. Group differences were assessed using Kruskal–Wallis tests, followed by Holm–Bonferroni-adjusted pairwise comparisons where applicable. Statistical significance is denoted by *P < 0.05, **P < 0.01, and ***P < 0.001. Exact P values are shown for selected comparisons. Recordings were obtained across three independent differentiation batches per cell line. Representative EPSC traces are displayed using standardized current-axis limits.

Analysis of spontaneous excitatory synaptic activity revealed significantly reduced EPSC frequency and amplitude in untreated T-derived DA neurons compared with controls (Figure 1f–j; frequency: n = 73 control and 51 patient neurons; P < 0.001; amplitude: n = 73 control and 47 patient neurons; P < 0.001). Isotretinoin treatment significantly increased EPSC frequency (n = 20 treated neurons; P < 0.01), whereas EPSC amplitude was not significantly altered (n = 20 treated neurons). Representative current traces were consistent with the reduced synaptic event frequency in untreated neurons and the partial recovery observed following treatment (Figure 1h–j).

At week 10, untreated T-derived DA neurons continued to exhibit significantly reduced excitability relative to controls, with decreases in both total and maximum evoked AP numbers (Figure 1k–o; n = 63 control and 28 patient neurons; both P < 0.001). Isotretinoin treatment significantly increased both measures compared with untreated patient-derived neurons (n = 22 treated neurons; both P < 0.001).

Despite these improvements in evoked firing, untreated patient-derived neurons continued to display significantly reduced EPSC frequency and amplitude compared with controls (Figure 1p–t; frequency: n = 83 control and 28 patient neurons; P < 0.01; amplitude: n = 76 control and 27 patient neurons; P < 0.05). Neither parameter was significantly restored by isotretinoin treatment (frequency: n = 22 treated neurons; amplitude: n = 21 treated neurons).

Together, these findings demonstrate persistent reductions in excitability and excitatory synaptic activity in T-derived DA neurons. Isotretinoin treatment increased evoked firing at both differentiation stages, whereas its effects on synaptic activity were more limited, suggesting differential responsiveness of intrinsic excitability and excitatory synaptic transmission.

#### Patient Z-derived DA neurons exhibit increased excitability and differentiation-stage-dependent responses to isotretinoin

In contrast to patient T, DA neurons derived from patient Z exhibited increased evoked neuronal firing relative to healthy controls.

At week 6, untreated Z-derived DA neurons displayed a modest but statistically significant increase in the total number of evoked APs compared with controls (Figure 2a–e; n = 71 control and 31 patient neurons; P < 0.05). Isotretinoin treatment significantly reduced total AP firing relative to untreated patient-derived neurons (n = 37 treated neurons; P < 0.05), shifting neuronal activity toward the control range. However, the maximum number of evoked APs did not differ significantly between untreated patient-derived neurons and controls (P = 0.18). Representative recordings illustrated the differences in evoked firing among control, untreated patient-derived, and treated patient-derived neurons (Figure 2c–e).

**Figure 2.**
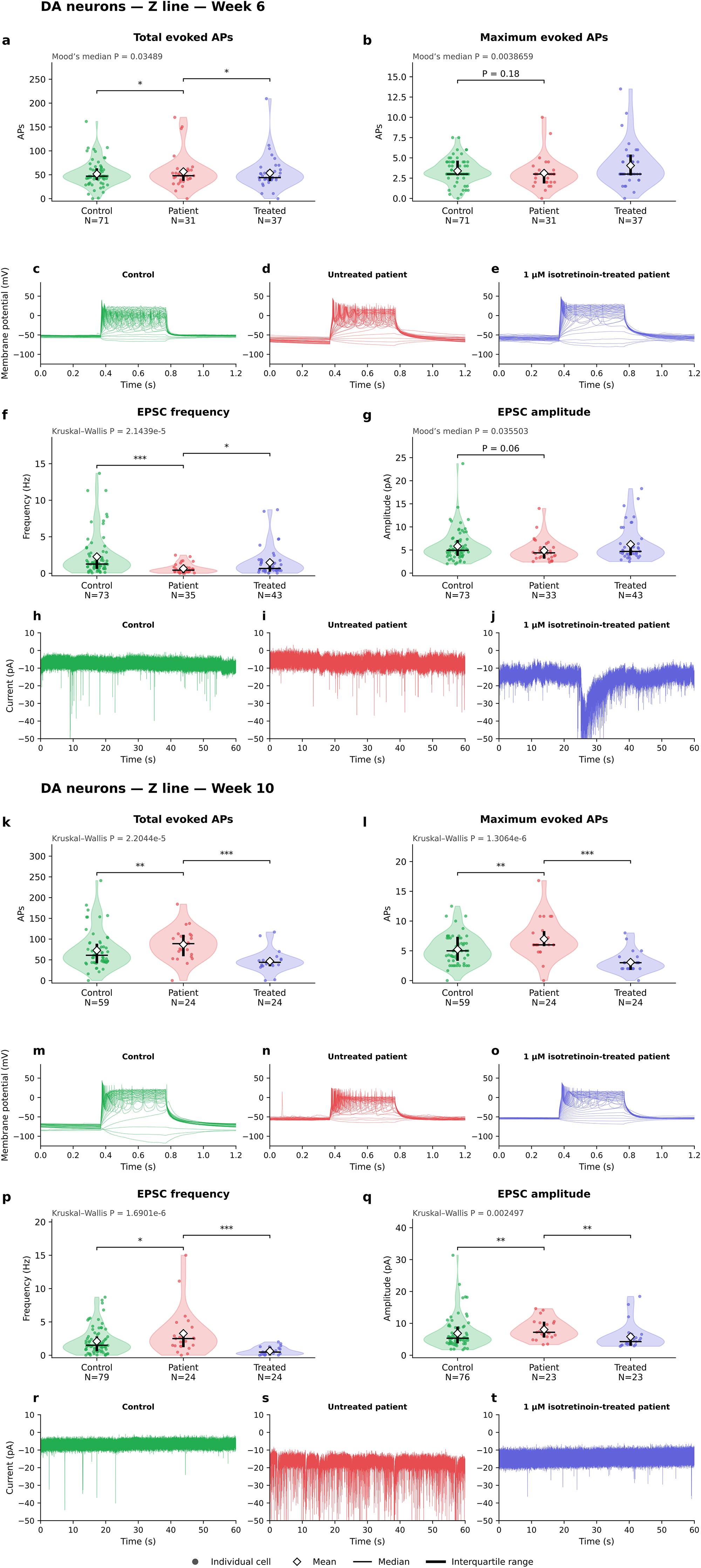
Electrophysiological characterization of SMS patient Z-derived DA neurons and the effects of isotretinoin treatment at weeks 6 and 10 of differentiation. Whole-cell patch-clamp recordings were performed in hiPSC-derived DA neurons from a healthy control and a patient with SMS (line Z). Patient-derived neurons were either untreated or exposed to 1 µM isotretinoin beginning on the first day of differentiation and maintained throughout neuronal differentiation. **Week 6 (a–j): (a, b)** Quantification of the total (a) and maximum (b) numbers of evoked APs in control (n = 71), untreated SMS (n = 31), and isotretinoin-treated SMS (n = 37) neurons. Untreated patient-derived neurons exhibited significantly increased total evoked AP numbers compared with controls (P < 0.05), whereas maximum AP numbers did not differ significantly (P = 0.18). Isotretinoin treatment significantly reduced total evoked AP numbers relative to untreated patient-derived neurons (P < 0.05), shifting this parameter toward control values. **(c–e)** Representative current-clamp recordings showing evoked AP responses to sequential depolarizing current injections in control (c), untreated SMS (d), and isotretinoin-treated SMS (e) neurons. **(f, g)** Quantification of EPSC frequency (f) and amplitude (g). EPSC frequency was assessed in control (n = 73), untreated SMS (n = 35), and isotretinoin-treated SMS (n = 43) neurons. Untreated patient-derived neurons exhibited significantly reduced EPSC frequency compared with controls (P < 0.001), whereas isotretinoin treatment significantly increased EPSC frequency relative to untreated patient-derived neurons (P < 0.05). EPSC amplitude was assessed in the same groups (n = 73, 33, and 43 neurons, respectively) and did not differ significantly between untreated patient-derived neurons and controls (P = 0.06). **(h–j)** Representative voltage-clamp recordings illustrating spontaneous EPSCs in control (h), untreated SMS (i), and isotretinoin-treated SMS (j) neurons. **Week 10 (k–t): (k, l)** Quantification of the total (k) and maximum (l) numbers of evoked APs in control (n = 59), untreated SMS (n = 24), and isotretinoin-treated SMS (n = 24) neurons. Untreated patient-derived neurons exhibited significantly increased total and maximum numbers of evoked APs compared with controls (both P < 0.01). Isotretinoin treatment significantly reduced both parameters relative to untreated patient-derived neurons (both P < 0.001), shifting evoked firing toward control values. **(m–o)** Representative current-clamp recordings showing evoked AP responses to sequential depolarizing current injections in control (m), untreated SMS (n), and isotretinoin-treated SMS (o) neurons. **(p, q)** Quantification of EPSC frequency (p) and amplitude (q). EPSC frequency was assessed in control (n = 79), untreated SMS (n = 24), and isotretinoin-treated SMS (n = 24) neurons; amplitude was assessed in 76, 23, and 23 neurons, respectively. Untreated patient-derived neurons exhibited significantly increased EPSC frequency (P < 0.05) and amplitude (P < 0.01) compared with controls. Isotretinoin treatment significantly reduced EPSC frequency (P < 0.001) and amplitude (P < 0.01) relative to untreated patient-derived neurons, shifting both parameters toward control values. **(r–t)** Representative voltage-clamp recordings illustrating spontaneous EPSCs in control (r), untreated SMS (s), and isotretinoin-treated SMS (t) neurons. Violin plots show distributions of individual neuronal measurements; points represent individual neurons, white diamonds indicate means, horizontal black lines indicate medians, and vertical black bars indicate interquartile ranges. Sample sizes (n) are indicated for each group. Group differences were assessed using Kruskal–Wallis tests, with Mood’s median tests where indicated, followed by Holm–Bonferroni-adjusted pairwise comparisons where applicable. Statistical significance is denoted by *P < 0.05, **P < 0.01, and ***P < 0.001. Exact P values are shown for selected comparisons. Recordings were obtained across three independent differentiation batches per cell line. Representative EPSC traces are displayed using standardized current-axis limits.

Despite the increased total evoked firing, untreated Z-derived DA neurons exhibited significantly reduced EPSC frequency compared with controls (Figure 2f–j; n = 73 control and 35 patient neurons; P < 0.001). Isotretinoin treatment significantly increased EPSC frequency relative to untreated patient-derived neurons (n = 43 treated neurons; P < 0.05), shifting this parameter toward control values. EPSC amplitude did not differ significantly between untreated patient-derived neurons and controls (n = 73 control, 33 patient, and 43 treated neurons; P = 0.06).

By week 10, untreated Z-derived DA neurons exhibited a more pronounced hyperexcitability phenotype, characterized by significantly elevated total and maximum evoked AP numbers compared with controls (Figure 2k–o; n = 59 control and 24 patient neurons; both P < 0.01). Isotretinoin treatment significantly reduced both parameters relative to untreated patient-derived neurons (n = 24 treated neurons; both P < 0.001), shifting evoked firing toward or below control levels.

At this later differentiation stage, untreated Z-derived neurons also exhibited significantly increased EPSC frequency (P < 0.05) and amplitude (P < 0.01) compared with controls (Figure 2p–t; frequency: n = 79 control and 24 patient neurons; amplitude: n = 76 control and 23 patient neurons). Isotretinoin treatment significantly reduced both EPSC frequency (P < 0.001) and amplitude (P < 0.01; frequency: n = 24 treated neurons; amplitude: n = 23 treated neurons), shifting these parameters toward or below the control range.

Collectively, these results reveal a distinct electrophysiological phenotype in Z-derived DA neurons, characterized by increased evoked firing and differentiation-stage-dependent changes in excitatory synaptic activity. Unlike the effects observed in patient T, isotretinoin reduced elevated excitability in patient Z, particularly at week 10, when treatment also reduced the increased EPSC frequency and amplitude.

#### Patient T-derived DG neurons exhibit reduced excitability and synaptic activity, with partial responses to isotretinoin

To determine whether the electrophysiological abnormalities observed in DA neurons extended to another neuronal population, whole-cell patch-clamp recordings were performed in iPSC-derived DG granule neurons at weeks 4 and 8 of differentiation.

At week 4, untreated DG neurons derived from patient T exhibited significantly reduced excitability compared with controls, as demonstrated by lower total and maximum evoked AP numbers (Figure 3a–e; n = 72 control and 47 patient neurons; both P < 0.001). Isotretinoin treatment significantly increased the total number of evoked APs relative to untreated patient-derived neurons (n = 58 treated neurons; P < 0.05), whereas maximum AP numbers were not significantly altered. Representative membrane potential recordings illustrated the reduced firing capacity of untreated patient-derived neurons and the treatment-associated increase in evoked activity (Figure 3c–e).

**Figure 3.**
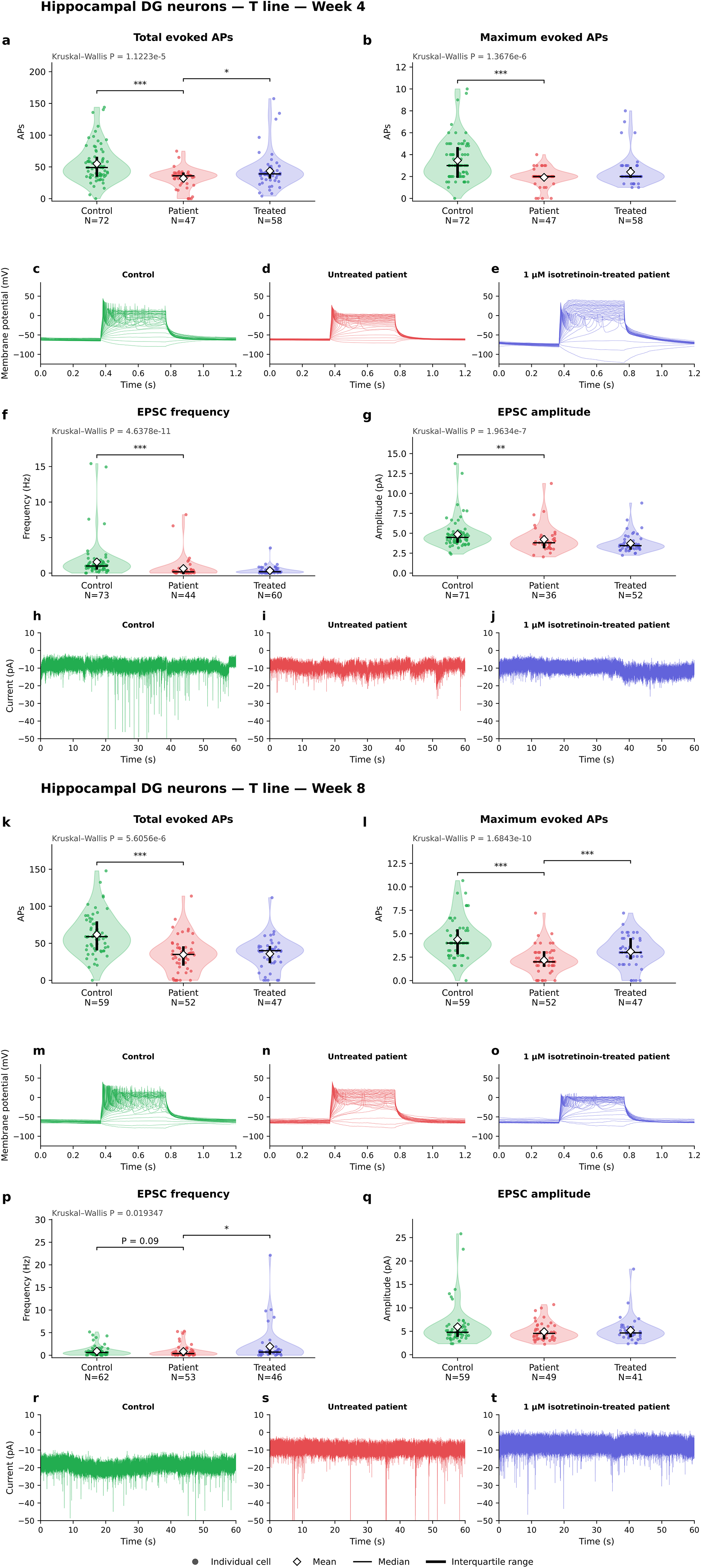
Electrophysiological characterization of SMS patient T-derived DG neurons and the effects of isotretinoin treatment at weeks 4 and 8 of differentiation. Whole-cell patch-clamp recordings were performed in hiPSC-derived DG neurons from a healthy control and a patient with SMS (line T). Patient-derived neurons were either untreated or exposed to 1 µM isotretinoin beginning on the first day of differentiation and maintained throughout neuronal differentiation. **Week 4 (a–j): (a, b)** Quantification of the total (a) and maximum (b) numbers of evoked APs in control (n = 72), untreated SMS (n = 47), and isotretinoin-treated SMS (n = 58) neurons. Untreated patient-derived neurons exhibited significantly reduced total and maximum numbers of evoked APs compared with controls (both P < 0.001). Isotretinoin treatment significantly increased the total number of evoked APs relative to untreated patient-derived neurons (P < 0.05), whereas no significant treatment-associated increase in maximum AP number was indicated. **(c–e)** Representative current-clamp recordings showing evoked AP responses to sequential depolarizing current injections in control (c), untreated SMS (d), and isotretinoin-treated SMS (e) neurons. **(f, g)** Quantification of EPSC frequency (f) and amplitude (g) in control, untreated SMS, and isotretinoin-treated SMS neurons (frequency: n = 73, 44, and 60; amplitude: n = 71, 36, and 52, respectively). Untreated patient-derived neurons exhibited significantly reduced EPSC frequency compared with controls (P < 0.001), without a significant treatment-associated improvement. EPSC amplitude was significantly lower in untreated patient-derived neurons than in controls (P < 0.01). **(h–j)** Representative voltage-clamp recordings illustrating spontaneous EPSCs in control (h), untreated SMS (i), and isotretinoin-treated SMS (j) neurons. **Week 8 (k–t): (k, l)** Quantification of the total (k) and maximum (l) numbers of evoked APs in control (n = 59), untreated SMS (n = 52), and isotretinoin-treated SMS (n = 47) neurons. Untreated patient-derived neurons exhibited significantly reduced total and maximum numbers of evoked APs compared with controls (both P < 0.001). Isotretinoin treatment significantly increased the maximum number of evoked APs relative to untreated patient-derived neurons (P < 0.001), whereas no significant treatment-associated increase in total AP number was indicated. **(m–o)** Representative current-clamp recordings showing evoked AP responses to sequential depolarizing current injections in control (m), untreated SMS (n), and isotretinoin-treated SMS (o) neurons. **(p, q)** Quantification of EPSC frequency (p) and amplitude (q) in control, untreated SMS, and isotretinoin-treated SMS neurons (frequency: n = 62, 53, and 46; amplitude: n = 59, 49, and 41, respectively). EPSC frequency did not differ significantly between untreated patient-derived neurons and controls (P = 0.09), but was significantly increased following isotretinoin treatment relative to untreated patient-derived neurons (P < 0.05). No significant differences in EPSC amplitude were indicated between the experimental groups. **(r–t)** Representative voltage-clamp recordings illustrating spontaneous EPSCs in control (r), untreated SMS (s), and isotretinoin-treated SMS (t) neurons. Violin plots show distributions of individual neuronal measurements; points represent individual neurons, white diamonds indicate means, horizontal black lines indicate medians, and vertical black bars indicate interquartile ranges. Sample sizes (n) are indicated for each group. Group differences were assessed using Kruskal–Wallis tests, followed by Holm–Bonferroni-adjusted pairwise comparisons where applicable. Statistical significance is denoted by *P < 0.05, **P < 0.01, and ***P < 0.001. Exact P values are shown for selected comparisons. Recordings were obtained across three independent differentiation batches per cell line. Representative EPSC traces are displayed using standardized current-axis limits.

Analysis of excitatory synaptic activity revealed significantly reduced EPSC frequency and amplitude in untreated T-derived DG neurons compared with controls (Figure 3f–j; frequency: n = 73 control and 44 patient neurons; P < 0.001; amplitude: n = 71 control and 36 patient neurons; P < 0.01). Neither parameter was significantly improved following isotretinoin treatment (frequency: n = 60 treated neurons; amplitude: n = 52 treated neurons).

At week 8, untreated T-derived DG neurons continued to exhibit significantly reduced total and maximum evoked AP numbers compared with controls (Figure 3k–o; n = 59 control and 52 patient neurons; both P < 0.001). Isotretinoin treatment significantly increased the maximum number of evoked APs relative to untreated patient-derived neurons (n = 47 treated neurons; P < 0.001), whereas total AP numbers were not significantly affected.

EPSC frequency showed a nonsignificant reduction in untreated patient-derived neurons compared with controls (Figure 3p–t; n = 62 control and 53 patient neurons; P = 0.09). However, isotretinoin treatment significantly increased EPSC frequency relative to untreated patient-derived neurons (n = 46 treated neurons; P < 0.05). No significant treatment-associated difference in EPSC amplitude was indicated (n = 59 control, 49 patient, and 41 treated neurons).

These findings demonstrate that T-derived DG neurons exhibit persistent reductions in intrinsic excitability, accompanied by alterations in excitatory synaptic activity. Isotretinoin increased selected electrophysiological parameters at both differentiation stages, although the responses were incomplete and varied between measures.

#### Patient Z-derived DG neurons exhibit persistent hyperexcitability with limited isotretinoin responsiveness

In contrast to patient T, DG neurons derived from patient Z exhibited increased excitability relative to healthy controls.

At week 4, untreated Z-derived DG neurons displayed significantly elevated total and maximum evoked AP numbers compared with controls (Figure 4a–e; n = 72 control and 46 patient neurons; total APs: P < 0.05; maximum APs: P < 0.001). Isotretinoin treatment did not significantly alter either parameter (n = 42 treated neurons). Representative membrane potential recordings illustrated the increased firing activity of untreated patient-derived neurons and the persistence of this phenotype following treatment (Figure 4c–e).

**Figure 4.**
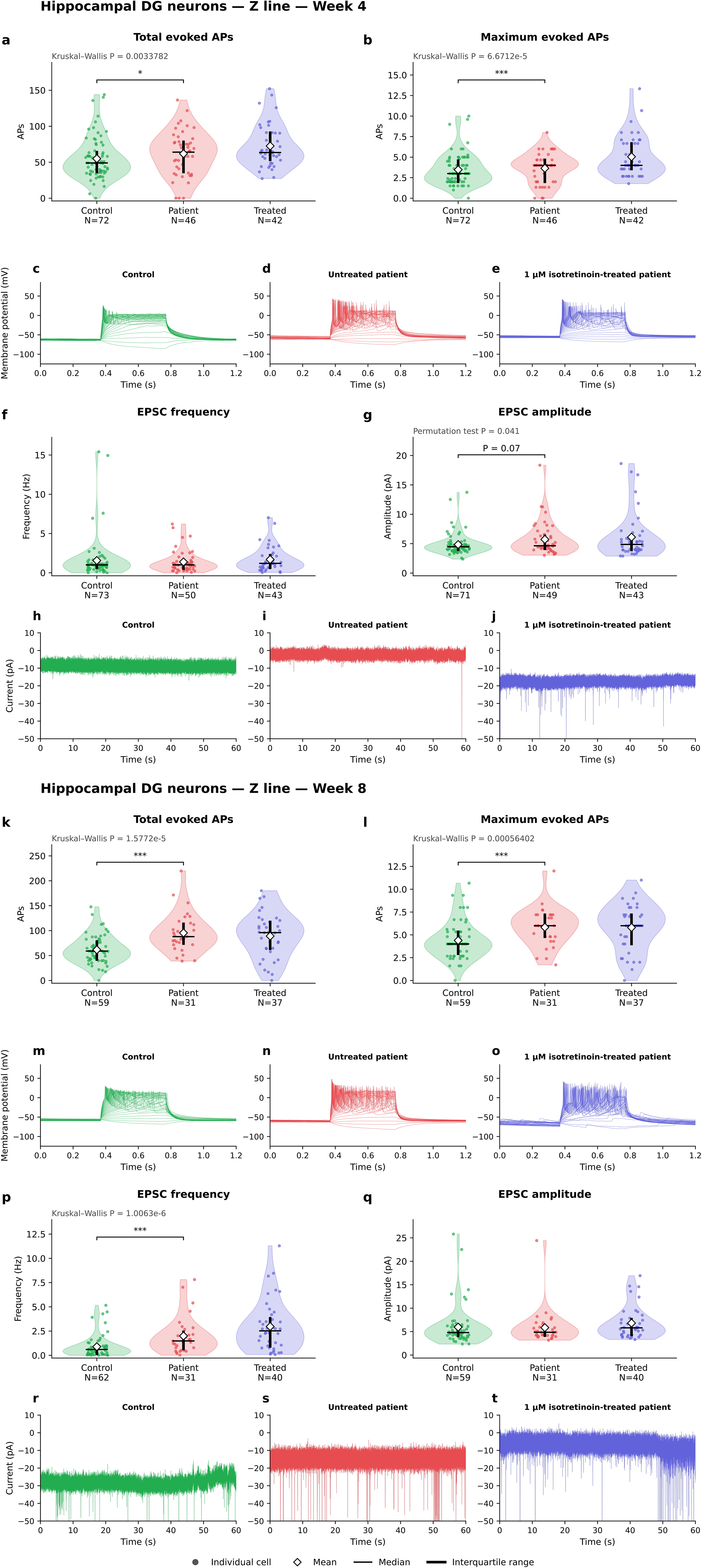
Electrophysiological characterization of SMS patient Z-derived DG neurons and the effects of isotretinoin treatment at weeks 4 and 8 of differentiation. Whole-cell patch-clamp recordings were performed in hiPSC-derived DG neurons from a healthy control and a patient with SMS (line Z). Patient-derived neurons were either untreated or exposed to 1 µM isotretinoin beginning on the first day of differentiation and maintained throughout neuronal differentiation. **Week 4 (a–j): (a, b)** Quantification of the total (a) and maximum (b) numbers of evoked APs in control (n = 72), untreated SMS (n = 46), and isotretinoin-treated SMS (n = 42) neurons. Untreated patient-derived neurons exhibited significantly increased total evoked AP numbers compared with controls (P < 0.05), as well as significantly increased maximum AP numbers (P < 0.001). Isotretinoin treatment did not significantly alter either parameter relative to untreated patient-derived neurons. **(c–e)** Representative current-clamp recordings showing evoked AP responses to sequential depolarizing current injections in control (c), untreated SMS (d), and isotretinoin-treated SMS (e) neurons. **(f, g)** Quantification of EPSC frequency (f) and amplitude (g). EPSC frequency was assessed in control (n = 73), untreated SMS (n = 50), and isotretinoin-treated SMS (n = 43) neurons. No significant differences in EPSC frequency were indicated between the experimental groups. EPSC amplitude was assessed in control (n = 71), untreated SMS (n = 49), and isotretinoin-treated SMS (n = 43) neurons. EPSC amplitude did not differ significantly between untreated patient-derived neurons and controls (P = 0.07), and no significant treatment-associated difference was indicated. **(h–j)** Representative voltage-clamp recordings illustrating spontaneous EPSCs in control (h), untreated SMS (i), and isotretinoin-treated SMS (j) neurons. **Week 8 (k–t): (k, l)** Quantification of the total (k) and maximum (l) numbers of evoked APs in control (n = 59), untreated SMS (n = 31), and isotretinoin-treated SMS (n = 37) neurons. Untreated patient-derived neurons exhibited significantly increased total and maximum numbers of evoked APs compared with controls (both P < 0.001). Isotretinoin treatment did not significantly alter either parameter relative to untreated patient-derived neurons. **(m–o)** Representative current-clamp recordings showing evoked AP responses to sequential depolarizing current injections in control (m), untreated SMS (n), and isotretinoin-treated SMS (o) neurons. **(p, q)** Quantification of EPSC frequency (p) and amplitude (q). EPSC frequency was assessed in control (n = 62), untreated SMS (n = 31), and isotretinoin-treated SMS (n = 40) neurons. Untreated patient-derived neurons exhibited significantly increased EPSC frequency compared with controls (P < 0.001), whereas isotretinoin treatment did not significantly alter EPSC frequency relative to untreated patient-derived neurons. EPSC amplitude was assessed in the same groups (n = 59, 31, and 40 neurons, respectively), with no significant differences indicated between experimental conditions. **(r–t)** Representative voltage-clamp recordings illustrating spontaneous EPSCs in control (r), untreated SMS (s), and isotretinoin-treated SMS (t) neurons. Violin plots show distributions of individual neuronal measurements; points represent individual neurons, white diamonds indicate means, horizontal black lines indicate medians, and vertical black bars indicate interquartile ranges. Sample sizes (n) are indicated for each group. Group differences were assessed using Kruskal–Wallis tests, with a permutation test where indicated, followed by Holm–Bonferroni-adjusted pairwise comparisons where applicable. Statistical significance is denoted by *P < 0.05, **P < 0.01, and ***P < 0.001. Exact P values are shown for selected comparisons. Recordings were obtained across three independent differentiation batches per cell line. Representative EPSC traces are displayed using standardized current-axis limits.

No significant differences in EPSC frequency were indicated among the experimental groups at week 4 (Figure 4f–j; n = 73 control, 50 patient, and 43 treated neurons). EPSC amplitude showed a nonsignificant difference between untreated patient-derived neurons and controls (n = 71 control, 49 patient, and 43 treated neurons; P = 0.07).

At week 8, untreated Z-derived DG neurons continued to exhibit increased excitability, with significantly elevated total and maximum evoked AP numbers compared with controls (Figure 4k–o; n = 59 control and 31 patient neurons; both P < 0.001). Isotretinoin treatment did not significantly alter either parameter (n = 37 treated neurons).

EPSC frequency was also significantly elevated in untreated patient-derived neurons compared with controls at week 8 (Figure 4p–t; n = 62 control and 31 patient neurons; P < 0.001). However, isotretinoin treatment did not produce a significant change in EPSC frequency (n = 40 treated neurons). EPSC amplitude did not differ significantly among the experimental groups (n = 59 control, 31 patient, and 40 treated neurons).

Together, these findings demonstrate persistent hyperexcitability in Z-derived DG neurons at both differentiation stages. Unlike Z-derived DA neurons, which exhibited substantial treatment-associated reductions in excitability and excitatory synaptic activity, Z-derived DG neurons showed limited electrophysiological responsiveness to isotretinoin.

#### Patient background and neuronal identity influence electrophysiological responses to isotretinoin

Collectively, electrophysiological characterization revealed contrasting functional phenotypes in SMS patient-derived neurons. Patient T exhibited predominantly reduced excitability and excitatory synaptic activity across DA and DG neuronal populations, whereas patient Z exhibited increased evoked firing, particularly at later differentiation stages.

Isotretinoin treatment was associated with differential functional responses depending on patient origin, neuronal identity, and differentiation stage. In T-derived neurons, treatment generally increased selected measures of evoked firing and excitatory synaptic activity, although the effects were incomplete. In Z-derived DA neurons, isotretinoin reduced elevated evoked firing and, at week 10, decreased the increased EPSC frequency and amplitude. In contrast, the elevated excitability of Z-derived DG neurons was not consistently modified by treatment.

These findings suggest that isotretinoin-associated electrophysiological modulation is not uniform across SMS-derived neuronal models and may depend on the baseline functional phenotype and cellular context.

#### Differential gene expression in SMS patient-derived DA neurons

Comparison of untreated patient Z-derived DA neurons with the familial control (ZC) and unrelated control (S) identified 3,598 and 6,484 DEGs, respectively (Figure 5b). Intersecting the two DEG sets and retaining concordant directions of change identified 1,897 consistently dysregulated genes (Figure 5a).

**Figure 5.**
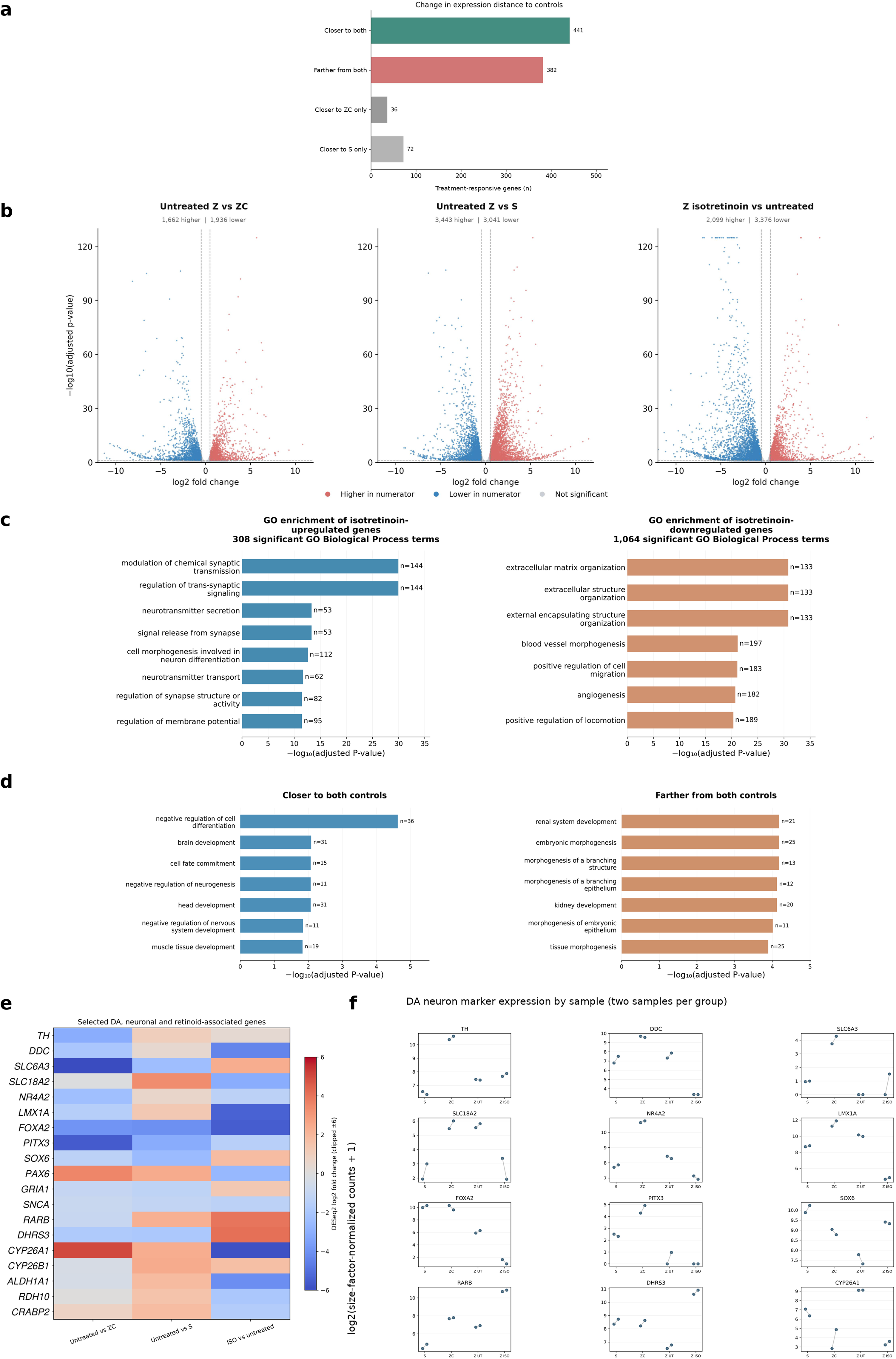
Transcriptomic alterations, functional enrichment, and isotretinoin-associated responses in SMS patient-derived DA neurons. (a) Classification of isotretinoin-responsive genes according to their transcriptional convergence toward or divergence from both mutation-negative control profiles. Of 1,897 genes consistently dysregulated in untreated patient-derived neurons relative to both controls, 931 were significantly altered following isotretinoin treatment. Among these, 441 genes (47.4%) shifted closer to both controls, 382 (41.0%) shifted farther from both, and 108 (11.6%) exhibited mixed responses, based on changes in absolute log fold differences. (b) Volcano plots illustrating differential gene expression in untreated patient-derived neurons versus the familial control (ZC; left), untreated patient-derived neurons versus the unrelated control (S; middle), and isotretinoin-treated versus untreated patient-derived neurons (right). The respective comparisons identified 3,598 (1,662 upregulated; 1,936 downregulated), 6,484 (3,443 upregulated; 3,041 downregulated), and 5,475 (2,099 upregulated; 3,376 downregulated) DEGs. Upregulation and downregulation are defined relative to the first condition in each comparison. (c) GO Biological Process enrichment analysis of genes upregulated and downregulated following isotretinoin treatment in SMS patient-derived DA neurons. Upregulated genes were significantly enriched in biological processes associated with neuronal communication and synaptic function, including modulation of chemical synaptic transmission, regulation of trans-synaptic signaling, neurotransmitter secretion, and regulation of membrane potential. In contrast, downregulated genes were predominantly enriched in processes associated with extracellular matrix organization, cell migration, and tissue morphogenesis. A total of 308 and 1,064 significantly enriched GO Biological Process terms were identified among upregulated and downregulated genes, respectively. Bars represent selected significantly enriched terms, with their lengths indicating −log of the Benjamini–Hochberg-adjusted *P*-value; gene counts (n) are annotated beside the bars. (d) GO Biological Process enrichment analysis of isotretinoin-responsive genes classified according to their transcriptional convergence toward or divergence from both mutation-negative control profiles. Enrichment was performed separately for genes moving closer to both controls (441 genes) and those moving farther from both controls (382 genes), identifying 17 and 119 significantly enriched GO terms, respectively. The ten most significantly enriched terms from each category are displayed. Bar labels represents the number of associated genes, and enrichment significance is expressed as −log of the Benjamini–Hochberg-adjusted p-value. (e) Heatmap illustrating log fold changes of selected genes involved in dopaminergic neuronal identity, synaptic function, and retinoid signaling. Comparisons include untreated SMS patient-derived neurons versus the familial control (ZC), untreated patient-derived neurons versus the unrelated control (S), and isotretinoin-treated versus untreated patient-derived neurons. Color intensity indicates the magnitude and direction of differential expression. (f) Normalized expression of selected dopaminergic and retinoid-associated genes in unrelated control (S), familial control (ZC), untreated SMS patient-derived (Z_UT), and isotretinoin-treated SMS patient-derived (Z_ISO) neurons. Expression values are presented as log (size-factor-normalized counts + 1). Individual points represent RNA-seq samples (n = 2 per group). DEGs were defined by a Benjamini–Hochberg-adjusted *p* < 0.05 and an absolute log fold change > 0.5. GO enrichment significance was determined using Benjamini–Hochberg-adjusted *p* < 0.05. RNA-seq samples originated from neuronal cultures generated within the same differentiation experiment; therefore, the analyses were considered exploratory.

Comparison of isotretinoin-treated and untreated patient-derived DA neurons identified 5,475 DEGs (Figure 5b).

To determine whether isotretinoin-associated transcriptional changes shifted patient-derived neurons toward the expression profiles of mutation-negative controls, a comparative analysis was performed using direct DESeq2 contrasts against both control lines.

For each eligible gene, the absolute log fold change relative to the familial and unrelated controls was compared with and without treatment. Among the 931 genes that were consistently dysregulated relative to both controls and responsive to isotretinoin, 441 (47.4%) moved closer to both control expression profiles and 382 (41.0%) moved farther from both. The remaining 108 genes (11.6%) showed divergent responses: 36 shifted toward the familial control (ZC) and away from the unrelated control (S), whereas 72 exhibited the opposite pattern (Figure 5a).

#### Isotretinoin treatment induces significant changes in synaptic pathways

GO Biological Process enrichment analysis revealed distinct functional enrichment profiles among genes upregulated and downregulated following isotretinoin treatment (Figure 5c). Isotretinoin-upregulated genes were significantly enriched in 308 GO Biological Process terms, with prominent enrichment of processes associated with neuronal communication, synaptic signaling, and neurotransmitter regulation. Selected significantly enriched terms included *modulation of chemical synaptic transmission* (144 genes, adjusted *P* = 1.14 × 10 ³ ), *regulation of trans-synaptic signaling* (144 genes, adjusted *P* = 1.14 × 10 ³ ), *neurotransmitter secretion* (53 genes, adjusted *P* = 4.72 × 10 ¹ ), and *regulation of membrane potential* (95 genes, adjusted *P* = 3.35 × 10 ¹²)(Figure 5c). Downregulated genes were enriched in 1,064 GO Biological Process terms, predominantly involving extracellular matrix organization, cell migration, and tissue morphogenesis (Figure 5c).

Together, these findings indicate that isotretinoin treatment induces extensive transcriptional remodeling involving synaptic communication, extracellular matrix regulation, and developmental signaling pathways.

Functional enrichment analysis of the 441 genes moving closer to both controls identified 17 significantly enriched GO Biological Process terms (Figure 5d). These included *negative regulation of cell differentiation* (36 genes, adjusted *P* = 2.4 × 10 ), *brain development* (31 genes, adjusted *P* = 0.0080), *cell fate commitment* (15 genes, adjusted *P* = 0.0083), and *negative regulation of neurogenesis* (11 genes, adjusted *P* = 0.0083). Genes moving farther from both controls yielded 119 significantly enriched GO Biological Process terms (Figure 5d).

These findings demonstrate that isotretinoin induces heterogeneous transcriptional responses. Although a substantial subset of initially dysregulated genes shifted toward control expression levels, a similarly large subset exhibited increased divergence. Genes moving toward control profiles were particularly associated with developmental and cell-fate regulatory processes, whereas genes moving farther from controls were enriched in morphogenetic and tissue-development pathways.

Given the established involvement of retinoid signaling in neuronal development and transcriptional regulation, genes associated with retinoic acid receptors, retinoid metabolism, and intracellular retinoid regulation were examined in greater detail (Figure 5e).

Isotretinoin treatment induced pronounced changes in retinoid-associated gene expression. The retinoic acid receptor gene *RARB* was strongly upregulated (log FC = 3.97, adjusted *P* = 1.68 × 10 ¹), together with *DHRS3* (log FC = 4.12, adjusted *P* = 4.81 × 10 ), which participates in retinoid metabolism. Expression of *RBP1*, encoding cellular retinol-binding protein 1, also increased substantially (log FC = 2.32, adjusted *P* = 7.32 × 10 ¹). In addition, *RARA* expression increased (log FC = 0.74, adjusted *P* = 4.80 × 10 ), whereas *RARG* expression decreased (log FC = −2.78, adjusted *P* = 2.58 × 10 ¹ ).

The expression of *RAI1* itself also increased following isotretinoin treatment (log FC = 0.477, adjusted *P* = 9.52 × 10 ), corresponding to an approximately 1.39-fold increase. Although statistically significant, this change did not meet the predefined absolute log fold-change threshold of 0.5 used for DEG classification.

Collectively, these findings demonstrate a pronounced isotretinoin-associated transcriptional response involving retinoic acid receptors, retinoid metabolism, and intracellular retinoid regulation. However, the direction of these changes was heterogeneous relative to the mutation-negative controls, indicating that activation of retinoid-responsive transcriptional programs was not uniformly associated with normalization of retinoid-related gene expression.

To further characterize isotretinoin-associated changes, genes involved in dopaminergic specification, neurotransmitter synthesis, vesicular transport, and synaptic function were examined. The heatmap summarizes differential-expression estimates (Figure 5e), while individual sample-level expression patterns are displayed separately (Figure 5f).

Several dopaminergic identity-associated genes were significantly downregulated following treatment, including LMX1A (log FC = −5.36), FOXA2 (−5.52), NR4A2 (−1.33), DDC (−4.40), and SLC18A2 (−3.07) (Figure 5e). In contrast, SOX6 increased (log FC = 1.81, adjusted P = 3.3 × 10 ¹ ), as did GRIA1 (log FC = 1.28, adjusted P = 8.7 × 10 ² ); both moved closer to the two control profiles (Figure 5F).

In contrast, TH, encoding tyrosine hydroxylase, exhibited a modest, nonsignificant increase (log FC = 0.36, adjusted P = 0.178). Similarly, SLC6A3, encoding the dopamine transporter, showed a positive estimated fold change that did not reach statistical significance (log FC = 2.36, adjusted P = 0.508). These estimated fold changes are summarized in Figure 5e; the corresponding sample-level expression patterns for these markers are shown in Figure 5f.

Conversely, SNCA, encoding α-synuclein, was significantly downregulated following treatment (log FC = −1.33, adjusted P = 4.93 × 10 ¹ ), with expression moving farther from both control profiles. The differential-expression estimate is shown in Figure 5e.

#### Summary of transcriptomic findings

Treatment-associated changes were enriched in pathways related to synaptic communication, extracellular matrix organization, neuronal development, and retinoid metabolism. Although 47.4% of the eligible disease-associated and treatment-responsive genes moved closer to both control expression profiles, 41.0% moved farther from both controls, demonstrating that isotretinoin-associated transcriptional remodelling was not uniformly restorative.

Substantial downregulation of several dopaminergic identity-associated genes suggested that isotretinoin exerted complex effects on neuronal transcriptional programs rather than uniformly enhancing dopaminergic maturation.

Together, these findings identify retinoid-responsive developmental, synaptic, and neuronal regulatory programs as prominent components of the isotretinoin response in SMS patient-derived DA neurons. Given that the DA RNA-seq analysis included a single SMS patient-derived line and samples originating from one differentiation batch, these findings should be considered exploratory and require validation in additional patient-derived lines and independent differentiation experiments.

#### Transcriptomic alterations in SMS patient-derived hippocampal neurons

To investigate the transcriptional consequences of *RAI1* deficiency in hippocampal neurons, bulk RNA sequencing was performed on iPSC-derived hippocampal neuronal cultures generated from two SMS patients (T and Z) and their respective familial controls (S and ZC). Patient-derived cultures were examined under untreated conditions and following exposure to 1 µM isotretinoin. Differential expression analyses incorporated family membership or patient-line identity as covariates, according to the experimental comparison.

PCA of variance-stabilized expression data revealed substantial transcriptomic variation across the hippocampal neuronal samples (Figure 6A). The first two principal components accounted for 60.6% and 18.2% of the total variance, respectively, indicating that the major sources of expression variability were captured by these components.

**Figure 6:**
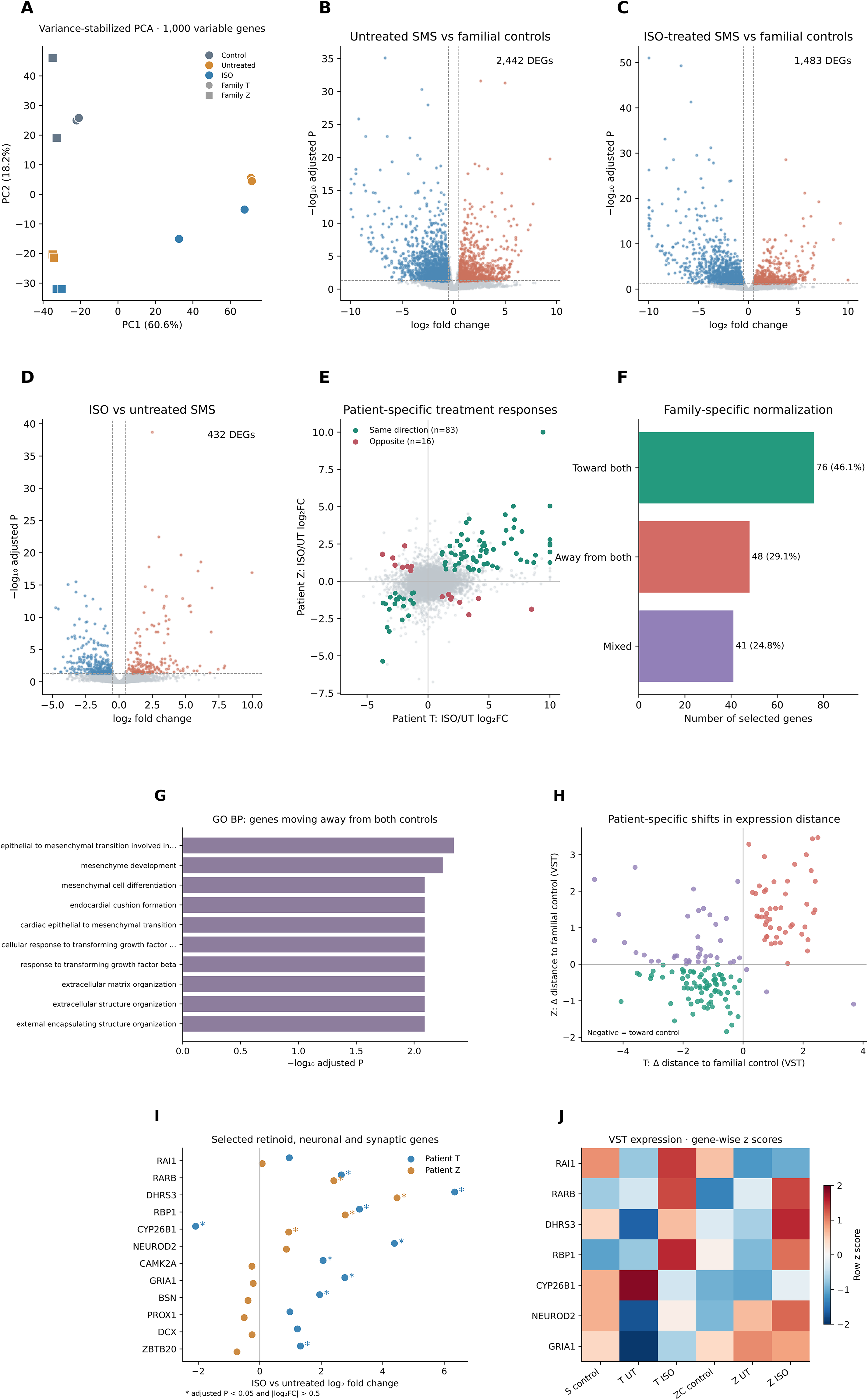
Transcriptomic alterations and isotretinoin-associated responses in SMS patient-derived hippocampal neurons. **(A)** PCA of variance-stabilized RNA-seq data from hippocampal neurons derived from two SMS patients (T and Z) and their respective familial controls (S and ZC), using the 1,000 most variable genes. PC1 and PC2 explained 60.6% and 18.2% of the total variance, respectively. **(B–D)** Volcano plots illustrating differential gene expression in untreated SMS-derived neurons versus familial controls (B), isotretinoin-treated SMS-derived neurons versus familial controls (C), and isotretinoin-treated versus untreated SMS-derived neurons (D). These comparisons identified 2,442, 1,483, and 432 DEGs, respectively. Patient-versus-control comparisons incorporated family membership as a covariate, whereas the combined treatment comparison incorporated patient-line identity. **(E)** Comparison of isotretinoin-associated log fold changes in patients T and Z. Among genes significantly altered in both patient lines, 83 exhibited concordant expression changes and 16 showed opposite directions. **(F)** Family-specific transcriptional convergence analysis. Among 165 genes significantly dysregulated in untreated SMS neurons and responsive to isotretinoin in the combined analyses, 76 moved toward both familial control profiles, 48 moved away from both, and 41 exhibited mixed responses. **(G)** Selected GO Biological Process terms enriched among genes moving away from both familial controls following isotretinoin treatment. No GO Biological Process terms reached the adjusted significance threshold among genes moving toward both controls. **(H)** Patient-specific changes in expression distance from the corresponding familial control, calculated using variance-stabilized expression values. Negative values indicate movement toward the familial control profile, whereas positive values indicate movement away. **(I)** Isotretinoin-associated log fold changes in selected genes involved in retinoid signaling, hippocampal neuronal development, and synaptic function, shown separately for patients T and Z. Asterisks indicate adjusted *P* < 0.05 and |log FC| > 0.5. **(J)** Heatmap illustrating selected gene-expression patterns across the two familial control lines and untreated and isotretinoin-treated SMS-derived hippocampal neurons. Variance-stabilized expression values are displayed as gene-wise standardized z-scores for the six cell-line/condition groups. For each hippocampal cell line, two RNA-seq samples per condition were collected from separate neuronal cultures originating from a single differentiation experiment. Isotretinoin (1µM) was administered exclusively to patient-derived neurons. DEGs were defined by Benjamini–Hochberg-adjusted *P* < 0.05 and |log FC| > 0.5. Differential-expression and transcriptional convergence analyses were interpreted as exploratory because independent differentiation experiments were not performed.

Comparisons between untreated SMS-derived neurons and familial controls identified 2,442 DEGs, including 911 upregulated and 1,531 downregulated genes (Figure 6B). Following isotretinoin treatment, the comparison between patient-derived neurons and familial controls identified 1,483 DEGs, comprising 454 upregulated and 1,029 downregulated genes (Figure 6C). Direct comparison of isotretinoin-treated and untreated SMS-derived neurons identified 432 treatment-associated DEGs, of which 171 were upregulated and 261 were downregulated (Figure 6D). All comparisons used a Benjamini–Hochberg-adjusted *P* value < 0.05 and an absolute log fold change > 0.5.

Although fewer genes differed significantly from familial controls following isotretinoin treatment, this reduction alone does not demonstrate transcriptional normalization, as differences in statistical significance may also reflect changes in expression variability and effect size.

Shared and patient-specific transcriptional responses to isotretinoin

To investigate interpatient variability in the transcriptional response to isotretinoin, differential expression was additionally assessed separately in hippocampal neurons derived from patients T and Z. These exploratory analyses identified 638 treatment-associated DEGs in patient T (344 upregulated and 294 downregulated) and 498 in patient Z (234 upregulated and 264 downregulated). Among genes meeting the differential-expression criteria in both patient lines, 83 exhibited concordant directions of change, whereas 16 displayed opposite responses (Figure 6E).

Examination of retinoid-associated genes revealed pronounced induction of *RARB*, *DHRS3*, and *RBP1* in both patient lines. *RARB* expression increased by log fold changes of 2.65 and 2.41 in T and Z, respectively, while *DHRS3* increased by 6.34 and 4.46 and *RBP1* by 3.25 and 2.78. All three genes met the differential-expression criteria in both patients, indicating a shared transcriptional response involving retinoid signaling and metabolism.

In contrast, *RAI1* expression did not change significantly following isotretinoin treatment in either patient line. Although patient T exhibited an estimated increase in *RAI1* expression (log fold change = 0.96), the change was not statistically significant after multiple-testing correction (adjusted *P* = 0.171). Patient Z exhibited a substantially smaller change (log fold change = 0.08; adjusted *P* = 0.999). These findings indicate that the observed retinoid-associated transcriptional responses occurred without a statistically confirmed increase in *RAI1* transcript abundance.

Additional differences were observed in genes associated with neuronal development and synaptic function. In patient T, isotretinoin significantly increased the expression of *NEUROD2* (log fold change = 4.38), *CAMK2A* (2.06), *GRIA1* (2.77), and *BSN* (1.95), whereas these genes did not meet the differential-expression criteria in patient Z. Furthermore, *CYP26B1*, a gene involved in retinoic acid metabolism, was significantly downregulated in T (log fold change = −2.09) but upregulated in Z (0.94). These observations suggest heterogeneity in downstream transcriptional responses despite shared induction of several retinoid-responsive genes (Figure 6I–J).

#### Isotretinoin-associated transcriptional convergence toward familial control profiles

To evaluate whether isotretinoin shifted disease-associated expression patterns toward those of familial controls, a family-specific expression-distance analysis was performed using variance-stabilized RNA-seq data. For each gene, the absolute difference between patient-derived and corresponding familial control expression was calculated before and after treatment. The analysis was restricted to genes significantly dysregulated in untreated SMS-derived neurons relative to controls and significantly responsive to isotretinoin in the combined differential-expression analysis.

A total of 165 genes fulfilled these criteria. Of these, 76 genes (46.1%) exhibited reduced expression distances from both familial controls following treatment, whereas 48 genes (29.1%) showed increased distances from both controls. The remaining 41 genes (24.8%) exhibited mixed responses, moving toward the familial control in one patient line but away in the other (Figure 6F,H).

Functional enrichment analysis of the 76 genes moving toward both controls did not identify any GO Biological Process terms meeting the adjusted significance threshold. In contrast, the 48 genes moving away from both controls yielded 56 significantly enriched GO Biological Process terms, including mesenchyme development, mesenchymal cell differentiation, extracellular matrix organization, and cellular responses to transforming growth factor-β (Figure 6G). Several enriched terms were associated with overlapping developmental and extracellular matrix-related gene sets.

Collectively, these results demonstrate that isotretinoin is associated with substantial transcriptional changes in SMS-derived hippocampal neurons, including shared induction of retinoid-associated genes and heterogeneous responses involving neuronal and synaptic pathways. Although a subset of treatment-responsive genes moved toward familial control expression profiles, transcriptional convergence was incomplete and varied between patient lines. These findings support further investigation of retinoid-associated transcriptional modulation in SMS-derived hippocampal neurons without establishing restoration of normal neuronal function.

#### Immune-associated transcriptional alterations differ between dopaminergic and hippocampal neuronal cultures

To examine immune-associated transcriptional alterations in SMS-derived neurons, genes involved in chemokine signaling, NF-κB-associated transcription, interferon responses, and complement regulation were assessed in the DA and hippocampal RNA-seq datasets. In untreated patient Z-derived DA neurons relative to familial control ZC, CCL2 (log FC = 1.87, adjusted P = 2.4 × 10 ²²), CXCL1 (2.93, 7.7 × 10 ² ), CXCL2 (2.43, 4.3 × 10 ), NFKB2 (1.51, 1.3 × 10 ¹²), RELB (1.63, 3.8 × 10 ), and TNFAIP3 (1.72, 1.6 × 10 ) were significantly upregulated. Increased expression was also observed for OAS3 (2.06, 2.8 × 10 ) and IFI44L (0.88, 0.0023). Baseline immune-associated expression patterns are shown in Figure 7A.

**Figure 7.**
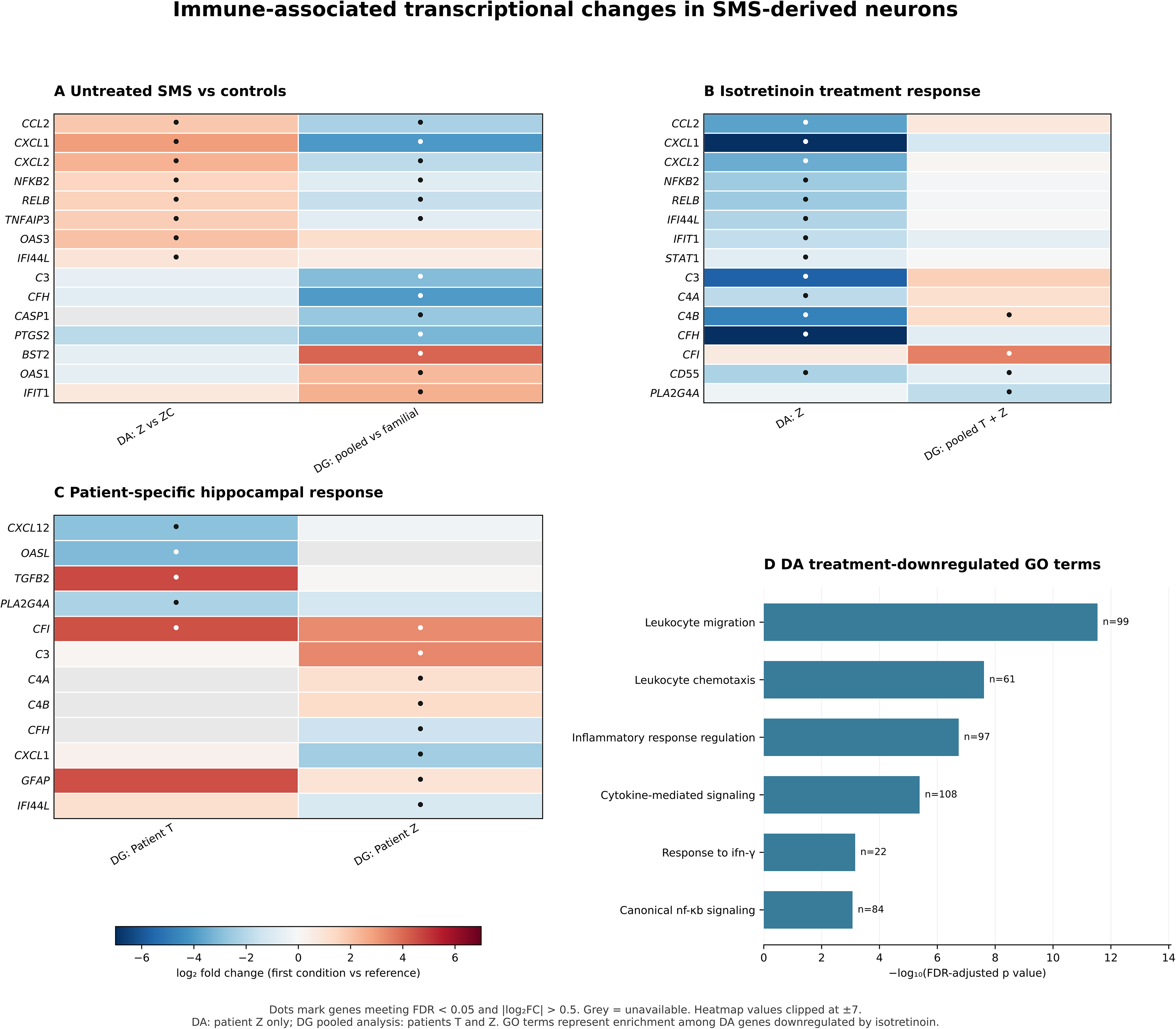
Differential modulation of immune-associated transcriptional programs by isotretinoin in SMS-derived dopaminergic and hippocampal neurons. (A) Heatmap showing selected immune- and inflammation-associated genes differentially expressed in untreated SMS-derived DA neurons from Patient Z relative to familial control ZC and in DG neurons relative to familial controls. (B) Treatment-associated changes in selected immune-related transcripts in DA neurons from Patient Z and in the combined hippocampal patient dataset. (C) Patient-specific transcriptional responses to isotretinoin in hippocampal neurons derived from Patients T and Z, highlighting differences in complement-associated, chemokine, and interferon-responsive gene expression. (D) Selected GO Biological Process terms significantly enriched among genes downregulated following isotretinoin treatment in DA neurons. Bar lengths represent −log -transformed Benjamini–Hochberg-adjusted p-values. Heatmap colors indicate log fold changes, with red representing increased expression and blue representing decreased expression. Dots indicate genes meeting the differential-expression thresholds of adjusted *p* < 0.05 and |log FC| > 0.5. Heatmap values are displayed on a scale capped at ±7. The hippocampal combined analysis includes both patient lines, whereas the DA transcriptomic analysis represents Patient Z. Enrichment of immune-associated GO terms indicates coordinated transcriptional changes but does not independently establish functional inflammatory activation.

Isotretinoin treatment of patient Z-derived DA neurons significantly reduced CXCL1 (log FC = −7.00, adjusted P = 3.0 × 10 ³ ), CCL2 (−3.76, 2.9 × 10 ²), CXCL2 (−3.49, 3.3 × 10 ), NFKB2 (−2.50, 4.5 × 10 ³¹), RELB (−2.55, 1.3 × 10 ), TNFAIP3 (−1.73, 8.2 × 10 ), and IFI44L (−2.10, 1.6 × 10 ¹ ). GO Biological Process analysis of genes downregulated after treatment identified enrichment of immune-related processes, including leukocyte migration, leukocyte chemotaxis, regulation of inflammatory response, and cytokine-mediated signaling (Figure 7D). These results indicate coordinated decreases in multiple immune-associated transcripts in the treated DA cultures (Figure 7B).

In the combined hippocampal comparison, untreated SMS-derived neurons showed lower expression of RELB (log FC = −1.61, adjusted P = 9.4 × 10 ¹ ), CASP1 (−2.63, 2.5 × 10 ), PTGS2 (−3.18, 1.4 × 10 ), and C3 (−3.01, 1.6 × 10 ) relative to familial controls. In contrast, interferon-responsive BST2 (4.08, 3.3 × 10 ) and OAS1 (2.25, 8.1 × 10 ) were upregulated. This mixed directionality differed from the pattern observed in untreated patient Z-derived DA cultures (Figure 7A).

The combined hippocampal treatment contrast identified increased CFI (log FC = 3.51, adjusted P = 5.7 × 10 ¹ ) and C4B (1.31, 0.016) (Figure 7B). Patient-specific analyses showed increased TGFB2 (4.61, 0.0039) and CFI (4.44, 0.030) in patient T, whereas patient Z exhibited increased CFI (3.28, 3.1 × 10⁻³⁴), C3 (3.35, 1.9 × 10⁻⁷), and C4B (1.28, 0.0035), accompanied by reduced CXCL1 (−2.44, 0.00089) and IFI44L (−1.06, 0.0027) (Figure 7C). Thus, immune-associated transcriptional responses to isotretinoin differed in their direction and gene composition between the DA and hippocampal datasets. These observations are descriptive and do not establish a statistically significant interaction between neuronal subtype and treatment.

## Discussion

### Patient-dependent electrophysiological abnormalities in SMS-derived neurons

A major finding of the present study was the identification of contrasting electrophysiological phenotypes in neurons derived from two patients with SMS. Despite the shared involvement of *RAI1* deficiency, neurons derived from patient T predominantly exhibited reduced excitability and excitatory synaptic activity, whereas neurons derived from patient Z displayed increased evoked firing, particularly at later differentiation stages. These differences were observed across DA and DG neuronal populations, suggesting that SMS-associated neuronal dysfunction is not characterized by a single electrophysiological phenotype.

The reduced excitability observed in T-derived neurons was evident at both differentiation stages and was accompanied by reduced EPSC frequency and, in several comparisons, amplitude. In contrast, Z-derived neurons exhibited increased evoked firing, with DA neurons developing more pronounced hyperexcitability and increased excitatory synaptic activity at later stages. Interestingly, Z-derived DA neurons exhibited increased evoked firing but reduced EPSC frequency at week 6, demonstrating that intrinsic excitability and excitatory synaptic transmission may be differentially affected during neuronal differentiation.

These findings are consistent with previous evidence implicating RAI1 in neuronal transcriptional regulation and synaptic homeostasis. Huang et al. (2016) demonstrated that RAI1 regulates genes involved in neuronal circuit assembly and communication, while Garay et al. (2020) identified a role for RAI1 in activity-dependent transcription and homeostatic synaptic scaling. Together, these studies support the possibility that disruption of RAI1-associated regulatory programs contributes to altered neuronal activity. Nevertheless, the opposing phenotypes observed in patients T and Z suggest that the consequences of RAI1 deficiency may depend on additional factors, including genetic background, mutation characteristics, neuronal differentiation state, and compensatory regulatory mechanisms.

### Isotretinoin differentially modulates neuronal excitability and synaptic activity

Isotretinoin treatment produced distinct electrophysiological responses depending on patient origin, neuronal population, and differentiation stage. In T-derived DA neurons, treatment significantly increased evoked firing at both weeks 6 and 10, partially counteracting the reduced excitability observed in untreated cultures. However, its effects on excitatory synaptic transmission were more limited, with increased EPSC frequency observed at week 6 but no significant restoration of EPSC frequency or amplitude at week 10.

In Z-derived DA neurons, isotretinoin produced an opposing response, reducing the elevated evoked firing observed in untreated cultures. At week 10, treatment also significantly decreased the increased EPSC frequency and amplitude, shifting these measures toward or below control values. These findings suggest that isotretinoin can modify neuronal activity in different directions depending on the underlying electrophysiological phenotype.

The hippocampal DG cultures demonstrated further differences in treatment responsiveness. In T-derived neurons, isotretinoin increased selected measures of evoked firing and EPSC frequency, although these effects varied with differentiation stage. Conversely, the increased excitability observed in Z-derived DG neurons persisted following treatment, indicating that isotretinoin did not consistently attenuate the electrophysiological abnormalities in this neuronal population.

Taken together, these observations suggest that isotretinoin does not exert a uniform excitatory or inhibitory effect on SMS-derived neurons. Instead, treatment-associated responses appear to depend on cellular context and baseline neuronal activity. Although several changes shifted electrophysiological parameters toward control values, these findings should be interpreted as partial functional modulation rather than complete restoration of neuronal physiology.

### Potential convergence between RAI1 and retinoid-dependent synaptic homeostasis

The electrophysiological responses observed in the present study may be considered in the context of established roles for RAI1 and retinoid signaling in synaptic homeostasis. Garay et al. (2020) demonstrated that RAI1 participates in activity-dependent transcriptional regulation and homeostatic synaptic scaling, suggesting that RAI1 deficiency may interfere with mechanisms responsible for maintaining stable neuronal activity.

Retinoic acid signaling has independently been implicated in homeostatic synaptic plasticity. Aoto et al. (2008) demonstrated that all-trans retinoic acid contributes to synaptic strengthening following prolonged suppression of neuronal activity through an RARα-dependent mechanism involving postsynaptic glutamate receptor regulation. Maghsoodi et al. (2008) further identified a role for retinoic acid in regulating local dendritic translation, providing a mechanism through which retinoid signaling may influence synaptic protein expression.

These findings provide a biological rationale for investigating whether retinoid-associated signaling can modify electrophysiological abnormalities associated with RAI1 deficiency. In the present study, isotretinoin increased evoked firing in neurons exhibiting reduced excitability and decreased firing in Z-derived DA neurons exhibiting elevated excitability. This pattern is compatible with context-dependent modulation of neuronal activity, although it does not establish engagement of homeostatic synaptic scaling.

Importantly, the mechanisms described in previous studies primarily involved all-trans retinoic acid, whereas the present experiments employed isotretinoin (13-cis-retinoic acid). Moreover, the current study did not directly assess synaptic scaling, AMPA receptor trafficking, or retinoid receptor activity. Consequently, the relationship between isotretinoin-associated electrophysiological changes and established retinoid-dependent homeostatic mechanisms remains hypothetical.

### Relationship between transcriptomic and electrophysiological responses

The electrophysiological and RNA-seq findings indicate that isotretinoin alters both neuronal activity and transcriptional programs in SMS-derived cultures, although the relationship between these responses remains unresolved. In patient Z-derived DA neurons, treatment was associated with differential expression of genes involved in synaptic signaling, neuronal differentiation, and retinoid metabolism. The same neuronal model also exhibited reduced evoked firing and excitatory synaptic activity at later electrophysiological assessment stages. Because RNA-seq and electrophysiology were performed at different differentiation time points and in separate cultures, these observations cannot establish a temporal or causal link between transcriptional and functional responses.

Enrichment of synaptic signaling pathways does not necessarily predict an increase in neuronal activity: isotretinoin-treated Z-derived DA neurons exhibited reduced firing and EPSC activity at week 10. Moreover, several dopaminergic identity-associated transcripts, including DDC, SLC18A2, NR4A2, LMX1A, and FOXA2, were reduced following treatment. These expression changes do not establish impaired neuronal maturation or loss of dopaminergic identity, because bulk RNA-seq cannot distinguish transcriptional regulation within individual neurons from differences in the proportions or differentiation states of cells in the cultures. Thus, treatment-associated electrophysiological modulation should not be equated with enhanced dopaminergic differentiation.

In hippocampal neurons, both patient lines exhibited induction of retinoid-responsive genes, including *RARB*, *DHRS3*, and *RBP1*. Nevertheless, the electrophysiological responses differed substantially. T-derived DG neurons showed treatment-associated increases in selected excitability and synaptic measures, whereas Z-derived DG neurons exhibited limited functional responsiveness despite evidence of retinoid-associated transcriptional activation.

This discrepancy suggests that engagement of retinoid-responsive transcriptional pathways does not necessarily translate into normalization of neuronal excitability. The induction of canonical retinoid-associated genes may therefore represent a shared molecular response to treatment, while downstream effects on neuronal physiology remain dependent on patient-specific and neuronal population-specific factors.

### Immune-associated transcriptional responses and neuronal context

An additional finding was the differential representation of immune-associated transcriptional programs in SMS-derived neuronal cultures. Untreated patient Z-derived DA neurons exhibited increased expression of several chemokines and NF-κB-associated transcripts, and isotretinoin reduced many of these transcripts. In hippocampal DG neurons, the combined analysis revealed a distinct pattern of immune-associated transcriptional alterations, characterized by increased expression of interferon-responsive genes (*BST2* and *OAS1*) alongside reduced expression of NF-κB-associated, inflammatory, and complement-related genes (*RELB*, *CASP1*, *PTGS2*, and *C3*). Following isotretinoin treatment, complement-associated transcriptional changes were particularly evident, with increased expression of *CFI* and *C4B* in the combined analysis. Patient-specific comparisons further revealed increased expression of *CFI*, *C3*, and *C4B* in patient Z-derived neurons, whereas patient T-derived neurons exhibited increased *CFI* and *TGF*β*2* expression.

These findings indicate context-dependent changes in immune-associated transcript abundance but do not establish neuroinflammation or anti-inflammatory activity. Bulk RNA-seq does not directly measure cytokine secretion, complement activation, or NF-κB signaling; differences in culture composition and neuronal maturation may also contribute. Protein-level and functional validation will be needed to determine whether these transcriptional changes reflect altered immune signaling.

### Influence of basal vitamin A supplementation and developmental timing on isotretinoin responses

An important consideration when interpreting the transcriptional and electrophysiological responses is the difference in basal retinoid supplementation between the neuronal differentiation protocols. DA neurons were differentiated in medium containing standard B27 Supplement from day 11, whereas hippocampal DG neurons were maintained with B27 Supplement minus Vitamin A throughout differentiation and maturation. Consequently, the two neuronal populations were exposed to different vitamin A supplementation conditions before and during isotretinoin treatment. Such differences may influence basal retinoid signaling, receptor responsiveness, and downstream transcriptional regulation.

The robust induction of retinoid-responsive genes, including *RARB*, *DHRS3*, and *RBP1*, in hippocampal neurons demonstrates that these cultures retained the capacity to respond transcriptionally to isotretinoin despite the absence of vitamin A supplementation through B27. This finding is particularly relevant because it suggests that isotretinoin-associated transcriptional responses can occur under conditions without intentional vitamin A supplementation through the B27 component. However, B27 minus Vitamin A does not necessarily establish a completely retinoid-free culture environment, as other medium constituents may contribute to basal retinoid availability.

Importantly, because neuronal identity and differentiation conditions differed simultaneously, the present study cannot determine whether differences in retinoid-associated responses between DA and hippocampal neurons were attributable to cell type, basal vitamin A availability, or other culture-specific factors. Future studies employing matched retinoid supplementation conditions and direct measurements of intracellular retinoid concentrations would help clarify these relationships.

The developmental timing of isotretinoin exposure represents an additional consideration. In the present study, isotretinoin was administered throughout neuronal differentiation, a period characterized by progressive changes in neuronal identity, intrinsic membrane properties, and synaptic connectivity. Retinoid signaling plays an established role in neuronal differentiation and maturation, suggesting that prolonged isotretinoin exposure may influence the establishment of neuronal functional properties rather than acting exclusively on mature neuronal activity.

This possibility is particularly relevant given the contrasting electrophysiological responses observed between patient-derived neuronal models. Isotretinoin increased evoked firing in the predominantly hypoexcitable neurons derived from patient T, whereas it reduced elevated excitability in Z-derived DA neurons. However, because treatment was maintained throughout differentiation, the present experiments cannot distinguish developmental effects from direct modulation of mature neuronal physiology. Future experiments involving acute isotretinoin administration or treatment initiated after neuronal maturation would help clarify the relative contributions of these mechanisms.

### Clinical observations and translational considerations

Importantly, electrophysiological responses were not uniformly restorative: in some T-derived DA measurements, treated values exceeded the control distribution, whereas other synaptic measures remained abnormal. In Z-derived DA neurons, treatment reduced hyperexcitability, in some comparisons shifting activity below control values, while Z-derived DG neurons showed limited responsiveness. Thus, the results support patient-, cell-type-, and maturation-dependent modulation rather than generalized normalization. Clinical interpretation is further limited by the single-patient observational design, concomitant levodopa/carbidopa and other possible confounders. The temporary isotretinoin dose increase was followed by mild liver-related laboratory abnormalities that reportedly normalized after dose reduction, underscoring the need for clinical safety monitoring. Biological samples used for iPSC generation were collected before the patient began isotretinoin, and the in vitro investigations commenced only after clinical treatment had begun; therefore, the experimental findings did not inform the initial prescribing decision. Controlled clinical and mechanistic studies would be needed to assess therapeutic relevance.

The longitudinal observations in patient T provide exploratory clinical context for the electrophysiological findings. Before isotretinoin initiation, progressive parkinsonian manifestations persisted despite partial benefit from levodopa/carbidopa and an earlier dose increase. Following the addition of isotretinoin during clinical care, neurological examinations documented gradual improvements in independent ambulation, balance, and selected functional motor abilities, as well as variable improvement in numerical tasks. Attention and conversational difficulties persisted. In vitro, isotretinoin increased evoked firing in T-derived hypoexcitable DA neurons at both examined stages, with a treatment-associated increase in EPSC frequency at week 6; selected measures in T-derived DG neurons also increased. These observations are directionally compatible with modulation of neuronal dysfunction, but the clinical and cellular outcomes are not directly equivalent and cannot be linked causally.

### Implications for SMS-associated neuronal dysfunction

The opposing functional phenotypes of T- and Z-derived neurons underscore the heterogeneity that can arise among SMS patient-derived models. The mother–child pair (ZC– Z) and sibling pair (S–T) provide family-matched comparisons, but these controls are not isogenic, and the two pairs differ in donor age, familial relationship, and potentially other background characteristics. With only two affected donors, the observed differences cannot be attributed specifically to mutation characteristics or generalized to all individuals with SMS.

Isotretinoin shifted some electrophysiological measures toward the respective control distributions but left others unchanged or shifted them beyond control values. Consequently, the present findings establish selective modulation in these models, not comprehensive restoration of disease-associated neuronal function.

The combination of electrophysiological and transcriptomic measurements is valuable because transcriptional shifts toward control profiles need not correspond to functional improvement. Conversely, changes in neuronal activity do not establish correction of RAI1-dependent molecular mechanisms. The convergence between RAI1-associated transcriptional regulation and retinoid-dependent homeostatic signaling remains a testable mechanistic hypothesis rather than a demonstrated pathway of action for isotretinoin.

### Study limitations

DA transcriptomic data were available for patient Z only, whereas the hippocampal analyses included both patient lines. Additional RNA sequencing of patient T-derived DA neurons is planned.

## Conclusions

In summary, patient-derived DA and hippocampal DG neurons revealed contrasting electrophysiological phenotypes in two individuals with SMS. Isotretinoin modified selected measures of excitability and synaptic activity in patient-, neuronal population-, and differentiation-stage-dependent directions, without uniformly restoring control-like function.

Transcriptomic analyses identified extensive retinoid-associated changes, including alterations in synaptic, developmental, and immune-associated gene programs. The incomplete correspondence between transcriptional and functional responses, together with the limited donor cohort and replication structure, supports further mechanistic investigation rather than a conclusion of disease correction or clinical efficacy.

## Competing Interests

The authors declare no competing interests.

## Funding

This research was supported the Zuckerman STEM leadership program, ISF 1437/26, and by the University of Haifa and the IBBRC.

## Data Availability

The datasets generated and analyzed during the current study, including bulk RNA-sequencing data, will be made available through an appropriate public repository upon publication. Additional data supporting the findings of this study may be obtained from the corresponding author upon reasonable request, subject to applicable ethical and privacy restrictions.

## Notes

### Competing Interest Statement

The authors have declared no competing interest.

